# Deciphering the Impact of Genetic Variation on Human Polyadenylation

**DOI:** 10.1101/2022.05.09.491198

**Authors:** Johannes Linder, Anshul Kundaje, Georg Seelig

## Abstract

Genetic variants that disrupt polyadenylation can cause or contribute to genetic disorders. Yet, due to the complex cis-regulation of polyadenylation, variant interpretation remains challenging. Here, we introduce a residual neural network model, *APARENT2*, that can infer 3’-cleavage and polyadenylation from DNA sequence more accurately than any previous model. This model generalizes to the case of alternative polyadenylation (APA) for a variable number of polyadenylation signals. We demonstrate APARENT2’s performance on several variant datasets, including functional reporter data and human 3’ aQTLs from GTEx. We apply neural network interpretation methods to gain insights into disrupted or protective higher-order features of polyadenylation. We fine-tune APARENT2 on human tissue-resolved transcriptomic data to elucidate tissue-specific variant effects. Finally, we perform in-silico saturation mutagenesis of all human polyadenylation signals and compare the predicted effects of >44 million variants against gnomAD. While loss-of-function variants were generally selected against, we also find specific clinical conditions linked to gain-of-function mutations. For example, using APARENT2’s predictions we detect an association between gain-of-function mutations in the 3’-end and Autism Spectrum Disorder.

## Introduction

Almost all human mRNA transcripts undergo cleavage and polyadenylation (pA). The position and efficiency of 3’ cleavage are controlled by a complex cis-regulatory code, the polyadenylation signal (PAS) (Figure 1A). The PAS consists of a core hexamer, typically AATAAA, and surrounding upstream and downstream sequence elements which together recruit the core processing machinery (CFIm, CstF, CPSF and hFIP1) [1, 2, 3, 4]. A large number of auxiliary factors, including hnRNP F/H/I, SRSF proteins, PABPC1, Ptbp2, HuR and Nova further modulate pA strength by binding to sequence motifs in the PAS [5, 6, 7, 8, 9, 10]. Adding an extra layer of complexity, the exact variants of these motifs, their relative positioning and interactions with structural motifs such as stem loops determine their cooperative, or antagonistic, effects [11]. Moreover, more than 70% of human genes contain multiple PASs (Alternative Polyadenylation, or APA), resulting in RNA isoforms with distinct 3’ ends (Figure 1B) [12, 13, 14]. The most common form of APA is the occurrence of two or more competing PASs in the 3’ untranslated region (3’ UTR) [1]. While all isoforms code for the same protein, their characteristics such as RNA stability or translation efficiency may vary considerably, as miRNA binding sites and other regulatory elements could have been removed from the shorter isoforms [15]. Less commonly, polyadenylation can also occur within introns, resulting in truncated protein isoforms.

**Figure 1:**
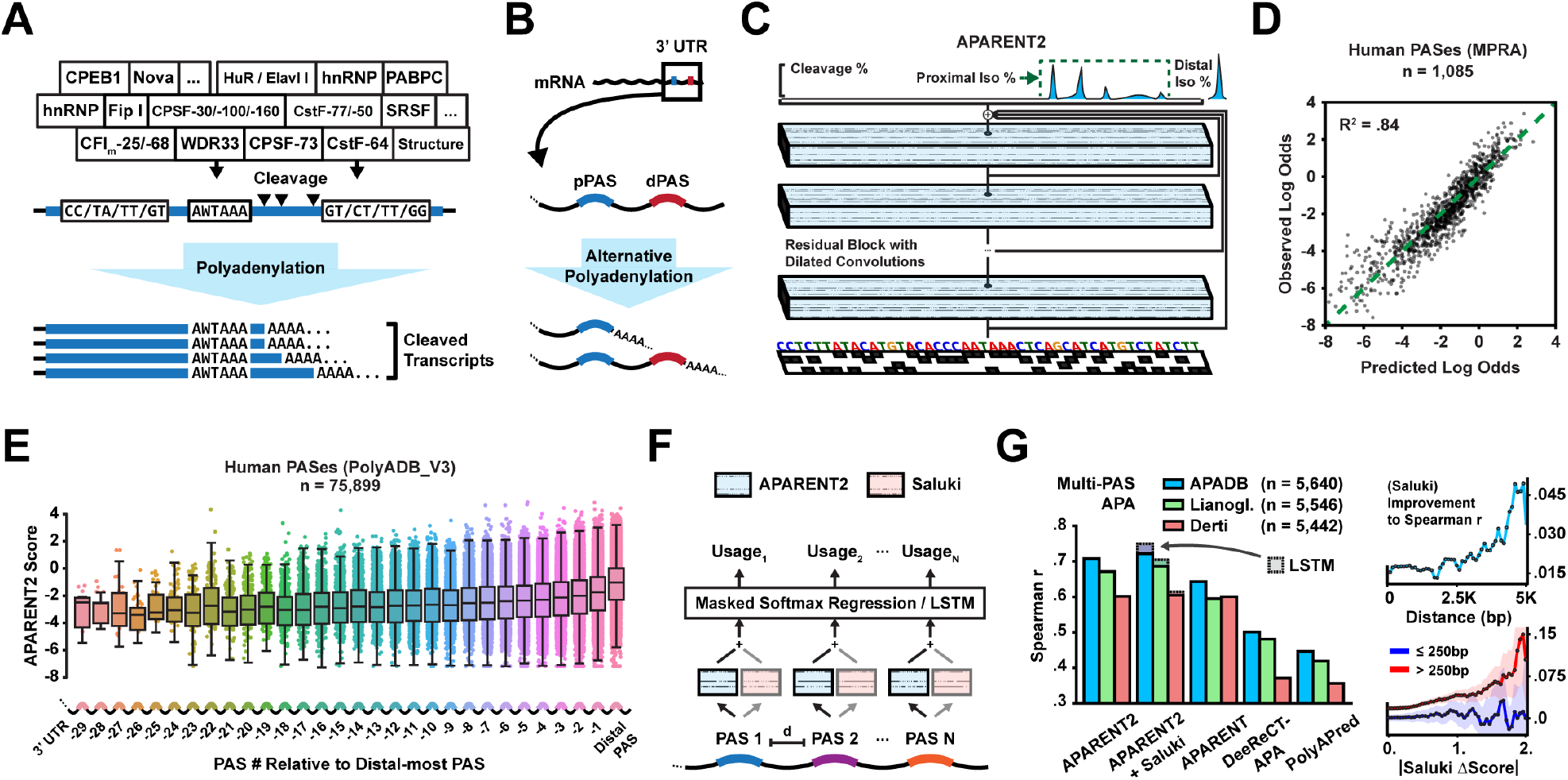
**A** Core processing elements, auxiliary RBPs and other determinants influence polyadenylation signal affinity. **B** Illustration of Tandem 3’ UTR Alternative Polyadenylation (APA) in pre-mRNA. **C** Residual neural network architecture. A one-hot coded representation of the PAS is used to predict the 3’ cleavage distribution. **D** Predicted vs measured proximal isoform log odds of native human 3’ UTR PASs measured in an MPRA (*n* = 1, 085). **E** Predicted logit score of all human PASs as a function of PAS # relative to the distal-most PAS. **F** Masked softmax regression (or a LSTM) for predicting multi-PAS isoform proportions given APARENT2 and Saluki scores as input. **G** Left: Comparison of correlation between predicted and measured distal isoform proportions from tissue-pooled native data (20-fold cross-validation). Each model predicts logit scores which are used to fit a multi-PAS regressor. LSTM performance shown as shaded bars. Right: Improvement in spearman r when using Saluki scores in addition to APARENT2 as input; (top) as a function of the min distance between at least one adjacent pair of PASs; (bottom) as a function of the min difference in Saluki score between at least one pair of PASs (blue / red = genes with PAS distances *≤*250*bp* / *>*250*bp*; shaded area = 90% confidence interval estimated by 10, 000-fold bootstrapping).

Assessing the impact of genetic variation on pA is important in both research and clinical settings, as several mutations that disrupt APA isoform abundances have been implicated in disease [16, 17, 18]. Even single PASs in 3’ UTRs without competing signals may have finely tuned functions, as weak mutations in such PASs can affect stability [19]. While genome-wide association studies (GWAS) and mapping of APA QTLs (3’ aQTLs) are powerful tools for finding statistical links between variants and phenotype [20, 21], they require a relatively large sample size and are less useful for rare or *de novo* variants [22, 23]. In a complementary approach, deep learning models that predict the *functional* impact of variants from sequence have been successful at classifying disruptive mutations, regardless of population frequency [24, 25, 26, 27, 28, 29, 30, 31]. Such sequence-predictive models have even been developed for pA [32, 33, 34, 35]. In particular, we previously trained a convolutional neural network (CNN) called APARENT for APA prediction [36].

Inspired by the recent success of deep residual networks applied to splicing and transcription factor binding prediction [27, 37], we here introduce *APARENT2*, a sequence-based residual neural network for 3’ cleavage prediction at base-pair resolution. We systematically compare the performance of APARENT2 to other models at the task of predicting disruptive variants, using functional MPRA data of 12, 350 single nucleotide variants (SNVs) from ClinVar and HGMD [36, 38, 39, 40] as well as scanning mutagenesis data from an assay of more than 12, 000 PASs [35]. We further compare the models on 366 high-confidence human 3’ aQTLs in 44 tissues from GTEx [20] and 58 aQTLs measured among 52 HapMap Yoruba human lymphoblastoid cell lines [21]. In all tests, APARENT2 significantly outperforms all state-of-the-art APA models. By combining APARENT2 with auxiliary tissue-specific models that we learn from native transcriptomic data in tissues and cell types expected to be differentially polyadenylated [41], we are able to provide residual variant predictions in testis, ovary, B-cell lymphocytes and brain that further boost performance on GTEx 3’ aQTLs.

In-silico interpretation methods have been applied extensively to assess the impact of genetic variants on the underlying cis-regulatory code [42, 37, 30, 43, 44, 45, 46]. Here, we use a mask-based interpretation method for neural networks – Scrambling – to elucidate higher-order features responsible for the predicted variant effects [47]. Specifically, we extend Scramblers to find the minimal set of features which explain the functional differences between a variant and wildtype sequence. With this approach, we discover super-additive interactions such as those between the CFIm-binding motif TGTA and AU-rich elements, or motifs that are differentially more active in brain and testis. We also find that some human PASs contain protective core hexamers (CSEs) that can initiate polyadenylation when the main CSE is disrupted. To understand the evolutionary constraints of polyadenylation in humans, we cross-reference the predicted effects of all potential 44 million polyadenylation SNVs against the 2.8 million PAS variants observed in gnomAD [48]. We find that loss-of-function variants occur *∼*2.5-fold less frequently in common variants (AF *>*10%) compared to singletons. However, when applying APARENT2 to a cohort study of Autism Spectrum Disorder (ASD), we found a *∼*3-fold enrichment of gain-of-function PAS mutations in cases (fisher’s exact p = 2.2*×*10^*−*4^) [49].

## Results

### A Residual Neural Network for Predicting 3’ Cleavage

Given recent advances in deep learning, we first asked whether an updated neural network architecture could improve on the performance of current state-of-the-art predictors such as APARENT. To this end, we trained a deep residual network on a re-processed version of the APA MPRA of Bogard et al. [36]. These data contain *>*3.3 million APA reporters with randomized proximal PAS sequence measured within 12 diverse 3’ UTR contexts. Briefly, the MPRA data was re-processed to map 3’ cleavage reads at basepair resolution for some missing UTR contexts (see Methods for details). The network, which is illustrated in Figure 1C and is referred to as APARENT2, is architecturally similar to SpliceAI [27] and BPNet [37]. Through a sequence of 28 Residual Blocks [50], each block consisting of two layers of dilated convolutions and a skip connection (Supplementary Figure S1A-B), the network transforms a one-hot coded representation of the input PAS (205 nt) into a predicted 3’ cleavage distribution. The last (206th) output of the network predicts the total isoform proportion of a far-away competing distal PAS (which in the training MPRA is non-random). For baseline comparisons, we also retrained a model with the original APARENT architecture on the re-processed version of the same MPRA (referred to as ConvNet below). To evaluate performance, we tested each network’s ability to infer total proximal isoform abundance on a set of 1, 085 native human PASs (also measured in the MPRA [36]) (Figure 1D). APARENT2 had significantly better correlation (*R*^2^ = 0.84) compared to the ConvNet baseline (*R*^2^ = 0.77; Supplementary Figure S1C). APARENT2 also had better correlation on held-out test data from the random MPRA (Supplementary Figure S1D).

Although APARENT2 was trained in the context of tandem APA, we note that the network effectively learns to score PASs relative to a fixed reference and we can thus interpret this score as an absolute measurement of PAS strength. Using APARENT2 as a PAS scoring function, we applied it to all human PASs in Polya DB V3 [51, 52]. In agreement with earlier analyses suggesting that distal signals are functionally more conserved [53], we found a near-perfect monotonically decreasing trend in predicted cis-regulatory strength as a function of PAS rank relative to the distal-most PAS of each gene (Figure 1E). The median strength of the proximal-most PAS was reduced *∼*6-fold compared to the distal-most PAS (Supplementary Figure S1E). We also successfully recapitulated binding motifs for several known pA mediators, including CFIm, CstF, HNRNPH2 and HuR by applying a motif discovery method, TF-MoDISco [54], to the APARENT2 predictions of 20, 000 PAS sequences from PolyA DB (Supplementary Figure S1F-G).

In the context of a multi-PAS gene, isoform abundance of a given PAS is determined not only by its intrinsic strength but also by the relative strength and distance of competing signals. Additionally, isoform abundances may be affected by the differential mRNA stability of the resulting 3’ UTR isoforms. To predict isoform proportions for genes with arbitrary numbers of PASs, we thus used native 3’-sequencing data to fit a multi-PAS regression model using the APARENT2 scores, the PAS distances, and the half-life of each isoform predicted by the Saluki model [55], as inputs (Figure 1F; spearman *r* ranged between 0.60 and 0.72 depending on data source when comparing measured to predicted distal isoform proportions with 20-fold cross-validation) [13, 41, 56]. When comparing to other APA models, including the CNN models PolyApredictor [35] and DeepPASTA [33] as well as the LSTM model DeeReCT-APA [34], APARENT2 was the most accurate at the task of multi-PAS prediction (Figure 1G) and pairwise PAS prediction (Supplementary Figure S1H-J). Switching the softmax regression layer of the multi-PAS model for a recurrent network (a LSTM [57]) resulted in only marginal performance gains (*r* increased by 0.01 to 0.028; Figure 1G).

While the overall improvement to predictive performance increased only modestly when including the Saluki half-life scores as input (spearman r increased by 0.015 on the APADB data), we noted that the improvement increased monotonically with larger differences between isoform lengths (Figure 1G, top right). For genes with large PAS distances (*>*250bp), a larger predicted difference in isoform stability was associated with larger improvement to predictive performance, while for genes with short isoforms (*≤*250bp) there was no improvement even for highly differentially stable transcripts (Figure 1G, bottom right). Taken together, these results suggest that APARENT2 can score cis-regulatory stability elements near the PAS, but that a more general stability model such as Saluki is beneficial for 3’ UTRs with long isoforms.

### Improved Prediction and Interpretation of Functional APA Variants

We next compared APARENT2 to APARENT, DeepPASTA, DeeReCT-APA and PolyApredictor at the tasks of classifying disruptive variants and estimating effect sizes (see Methods for details on how each model was used). We first analyzed our own variant MPRA [36], consisting of 12, 350 SNVs occuring near PASs of disease-implicated 3’ UTRs from ClinVar, HGMD or ACMG genes [38, 39, 40]. Figure 2A shows that the wildtype- and variant cleavage distributions predicted by APARENT2 match the measured peaks better than the original APARENT model. When comparing all models based on how well they could predict isoform fold changes and classify disruptive variants (|Fold Change| *>* 2), we found that APARENT2 had the highest overall accuracy (Figure 2B, Supplementary Figure S2A-B; Average Precision = 0.67; *R*^2^ = 0.69; *n* = 12, 350). Importantly, the performance gap of APARENT2 increased when looking only at a more challenging class of variants outside of the CSE.

**Figure 2:**
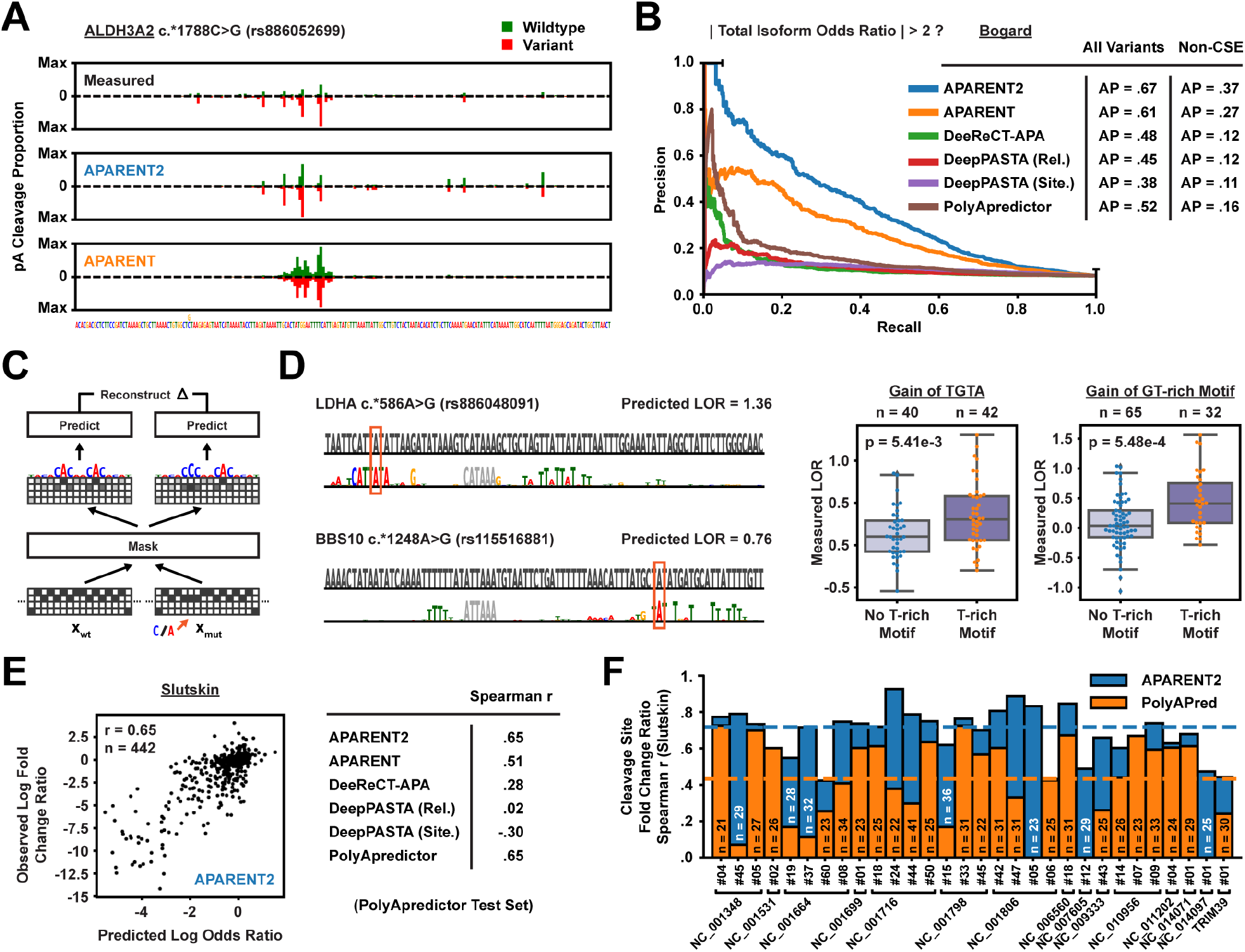
**A** Variant of uncertain significance from ClinVar (rs886052699) measured in the MPRA of Bogard et al. [36]. Shown are the measured and predicted 3’ cleavage distributions across the PAS. Green: Wildtype cleavage, Red: Variant cleavage. **B** Comparison of precision-recall curves when tasking each APA model with classifying disruptive APA variants (|fold change| *>* 2) from the MPRA of Bogard et al. [36] (n = 12, 350). The curves are shown for non-CSE variants only. **C** Mask-based variant interpretation, reconstructing the relative odds ratio between the wildtype and mutated sequence. **D** Interpretation of two ClinVar SNVs. Boxplots show measured LORs from the MPRA of Bogard et al. [36]. P-values are computed with two-sided t-tests. **E** Comparison of predicted vs measured RNA/DNA log fold change ratios on the data from Slutskin et al. [35] (n = 442). **F** Comparison of predicted vs measured RNA/DNA log fold change ratios at individual cleavage sites.

Given the increased performance of a more complex network architecture, we wanted to understand the types of higher-order regulatory features learned by APARENT2 that impact variant effect predictions. To this end, we used a neural network attribution method recently developed by our group – Scrambling – to detect contextual features responsible for the observed variant effects [47]. To interpret a mutation, we optimize a discretized attention mask to highlight a shared set of features (nucleotides) in the wildtype-and variant sequences that allows reconstruction of their predicted odds ratio (Figure 2C; see Methods for details).

In Figure 2D (and Supplementary Figure S2C) we interpret two gain-of-function variants, (1) rs886048091, which creates an upstream CFIm-binding motif (TGTA) and (2) rs115516881, which creates a downstream CstF-binding motif (GT-rich). These variants were predicted and measured to have variant fold changes significantly higher than the median fold change observed for other TGTA- or GT-creating mutations. Our interpretations elucidate cooperative interactions with downstream T-rich motifs, which explain the amplified variant effects. We also find support in the MPRA data of Bogard et al. [36], as T-rich elements in the DSE are associated with higher-amplitude TGTA- or GT-creating mutations (*p* = 5.41 *×* 10^*−*3^ and *p* = 5.48 *×* 10^*−*4^ respectively). Additionally, rs886048091 stabilizes the RNA secondary structure of the PAS (Supplementary Figure S2D) and the interpretation highlights altered base-pairing positions near the mutation. In Supplementary Figure S2E, we find that well-positioned T-rich elements are crucial determinants for *de novo* cleavage for mutations that create up- or downstream competing CSE hexamers.

### Predicting the Impact of Variants on Polyadenylation Signal Processing Efficiency

We further compared the models on a separate 3’ UTR MPRA which measured expression levels as a proxy for polyadenylation processing efficiency [35]. In this assay, a single PAS was inserted in each gene and RNA levels were found to vary over almost an order of magnitude with PAS strength and thus 3’-end processing efficiency. These data allow us to test our ability to infer intrinsic PAS strength independent of the presence of APA. We tested the models on a subset of the MPRA, which contains scanning mutagenesis measurements of several native PASs, including 572 viral PASs. We first compared the models on how well their predicted variant fold changes correlated with measured total RNA/DNA fold change ratios (Figure 2E, Supplementary Figure S2F). Here, APARENT2 and PolyApredictor, a model trained directly on these data, have identical correlations (spearman *r* = 0.65, *n* = 442), which is significantly higher than other models. However, when predicting variant fold change ratios at individual cleavage sites, APARENT2 was more accurate (Figure 2F, Supplementary Figure S2G; median spearman *r* = 0.72, total *n* = 1, 217).

### Functional Variant Predictions Correlate with Human APA QTLs

To assess the APA models on variant prediction within a native genomic context, we downloaded the recently published atlas of APA QTLs (3’ aQTLs) from GTEx v7 [20]. The majority of aQTL measurements involve distant SNPs far away from any PAS, which is beyond the scope of APARENT2. We thus narrowed the data to the subset of variants that occur close enough to the core hexamer of an annotated PAS in PolyA DB (within 50nt; n = 2, 043) (Figure 3A, Supplementary Figure S3A). We further filtered the data on lead SNPs (most significant SNP for a given APA event), resulting in a total of 366 GTEx 3’ aQTLs measured among 44 tissue types. We then tasked each model with inferring the aQTL effect size due to each variant (Figure 3A). APARENT2 had the highest median correlation across all tissues (spearman r = 0.61) and was followed by DeeReCT-APA (r = 0.48). APARENT2’s predictions correlated stronger with the aQTL effect sizes when using an increasingly larger p-value cutoff (Supplementary Figure S3B). We further benchmarked the models on a separate 3’ aQTL dataset [21], consisting of 58 SNVs occurring near annotated PASs among 52 HapMap Yoruba human lymphoblastoid cell lines (Figure 3B, Supplementary Figure S3C). APARENT2 again were the most correlated with the measured aQTLs (spearman r = 0.70). Finally, by comparing our variant predictions to 1, 007 intronic GTEx eQTLs and 2, 225 3’ UTR eQTLs [58], we validated an observation made by Mittleman et al. [21] that mRNA expression is significantly downregulated due to gain-of-function mutations in intronic PASs, possibly due to aberrant transcript truncation (Supplementary Figure S3D; across all GTEx tissues, we found that variant effects predicted by APARENT2 in weak intronic polyadenylation sites had a median negative correlation of *r* = *−*0.3 to the measured eQTL effect sizes).

**Figure 3:**
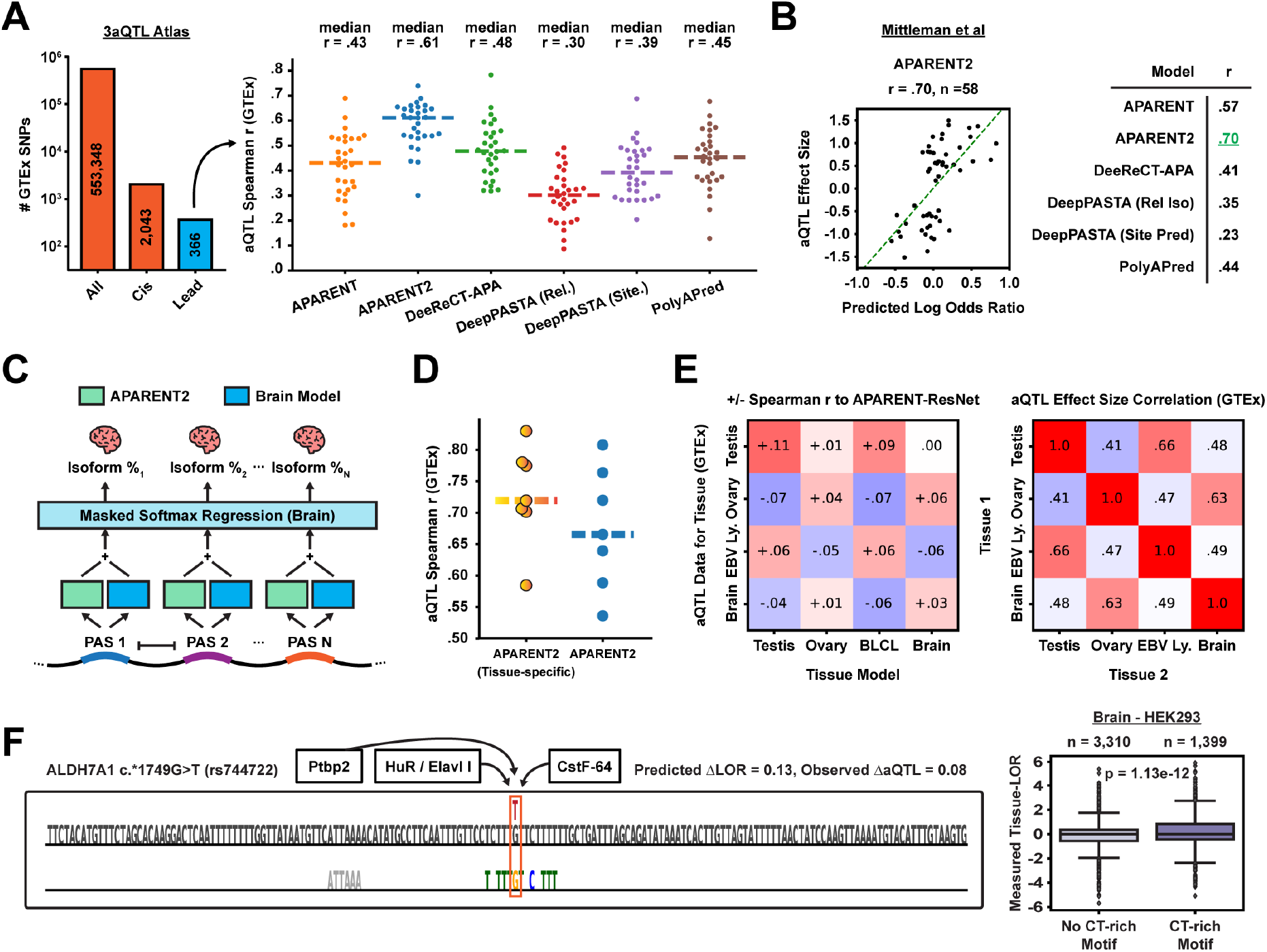
**A** Predicted vs measured GTEx 3’ aQTL effect size spearman r’s. Each dot corresponds to the spearman r in a particular tissue type. Left: Number of 3’ aQTLs, cis-acting 3’ aQTLs and lead 3’ aQTLs respectively. **B** Predicted vs estimated aQTL effect sizes of the Mittleman et al. [21] data (*n* = 58). **C** Multiple softmax regression for predicting tissue-specific PAS usage. APARENT2 (green) and the Tissue-model (blue) are used to score the strength of each PAS. **D** Predicted vs measured GTEx 3’ aQTL correlation for 7 tissue types (testis, ovary, B-cell lymphocytes, and 4 brain tissues), where the APARENT2 predictions have been scaled by the corresponding tissue model. **E** Left: Increase (red) or decrease (blue) in spearman r when using a particular tissue model to scale the 3’ aQTL predictions in a given GTEx tissue type. Right: Correlation of GTEx 3’ aQTL effect sizes between tissues. **F** Reconstructive mask for a GTEx SNP in the ALDH7A1 gene, with a tissue-specific effect in brain. Boxplot shows differential PAS usage in data from Lianoglou et al. [41].

### Tissue-specific Variant Prediction as Residual Learning

The 3’ aQTL effect sizes above are tissue-specific, yet APARENT2’s predictions are not. The reason we observe high correlation is because APA, for most genes and PASs, is not differentially regulated [59]. Thus, predictions of APARENT2, which was trained on MPRA data from HEK293 cells, correlate quite well across all tissues. However, we asked whether we could improve variant predictions on some of the aQTLs by combining native, tissue-specific 3’-end sequencing data with the single-cell line MPRA data in a hybrid model. Here we draw inspiration from earlier work by Cheng et al. [60], where tissue-specific splicing models were used to scale the variant predictions of a non-tissue specific model. This hybrid approach is motivated by the idea that the non-tissue specific model, which has been trained on a large MPRA, can provide more accurate baseline predictions. The tissue-specific models, then, are used only to predict residual up- or down-regulation due to tissue-specific *trans*-acting regulators and their cognate cis-acting motifs.

We focused on 4 human tissues and cell types that have previously been reported to exhibit differential polyadenylation [59]: testis, ovary, B-cell lymphocytes (BLCL) and brain. We downloaded publicly available 3’-end sequencing data for HEK293, testis, ovary, BLCL and brain [41] and mapped the RNA-Seq reads to annotated PASs in APADB [56]. In total, we collected APA isoform data for 6, 440 genes, each gene having between 2 and 10 PASs. First, in agreement with earlier studies suggesting that weaker PASs are upregulated in testis, we observed that the APARENT2 PAS score itself is predictive of differential usage in testis; the isoform odds ratio increases *∼*1.5-fold in testis for proximal PASs with scores *<*0 and distal competing signals with scores *>*0 (*p* = 2.5*×*10^*−*50^; Supplementary Figure S3E). Next, using these data, we trained four tissue-specific models to learn the residual APA regulation necessary to predict tissue-specific differences superimposed on the baseline APARENT2 predictions (Figure 3C; spearman r = 0.20 - 0.41 on held-out test data, Supplementary Figure S3F). After training, we used each tissue-specific model to scale the GTEx effect size predictions (Supplementary Figure S3G). With this approach, we raised the median aQTL spearman correlation from 0.66 to 0.72 (Figure 3D). Additionally, we found that the testis-specific model could be used to scale the B-cell GTEx predictions and vice versa, and similarly the Ovary- and Brain-specific models could be used interchangeably (Figure 3E; left, Supplementary Figure S3H). This relationship is consistent with the aQTL measurements from the GTEx atlas (Figure 3E; right).

Finally, we applied our mask-based attribution framework to interpret tissue-specific variants on the basis of the residual tissue models. In Figure 3F, we investigate GTEx SNP rs744722, which has a positive 3’ aQTL effect size in brain but a median negative effect size in other tissues. Our interpretation suggests that the variant modifies a T/GT/CT-rich motif by removing one of the Gs. We hypothesized that this SNP alters the affinity for CstF binding, which has an overall negative impact in most tissues, but has a net-positive effect in Brain due to the upregulated levels of HuR / Elavl I and Ptbp2, which are RBPs known to compete with CstF binding in T/GT/CT-rich regions [6, 7]. CLIP-data supports CstF binding overlapping the mutation site in ADH7A1 [61] and we find in the native transcriptomic training data that CT-rich motifs are associated with upregulated PAS usage in brain (*p* = 1.13*×*10^*−*12^). In Supplementary Figure S3I we interpret a similar loss-of-CstF binding mutation, which is observed to have a more negative effect size in testis compared to other tissues. Consistent with earlier studies, we find evidence that GT-rich motifs are associated with differential APA in Testis, which is likely due to elevated levels of CstF [62, 63, 64].

### Silent Hexamer Mutations are Protected by Functional Redundancy

The core cis-regulatory polyadenylation element in humans is the CSE hexamer motif, which in its canonical form is either AATAAA or ATTAAA but weaker nucleotide variants exist (4A) [1]. Reporter experiments measuring polyadenylation efficiency have recently shown that clinically benign CSE mutations often have lower functional effect sizes than expected [65]. To investigate this phenomenon at a larger scale, we collected all measured CSE mutations from the MPRA of Bogard et al. [36] (n = 628) and compared APARENT2’s variant effect predictions to the measurements (Figure 4B, left). APARENT2 can regress the effect sizes accurately (spearman r = 0.71) and the predictions generally separate the benign from pathogenic labels in ClinVar depending on whether a mutation is loss-of-function or silent.

**Figure 4:**
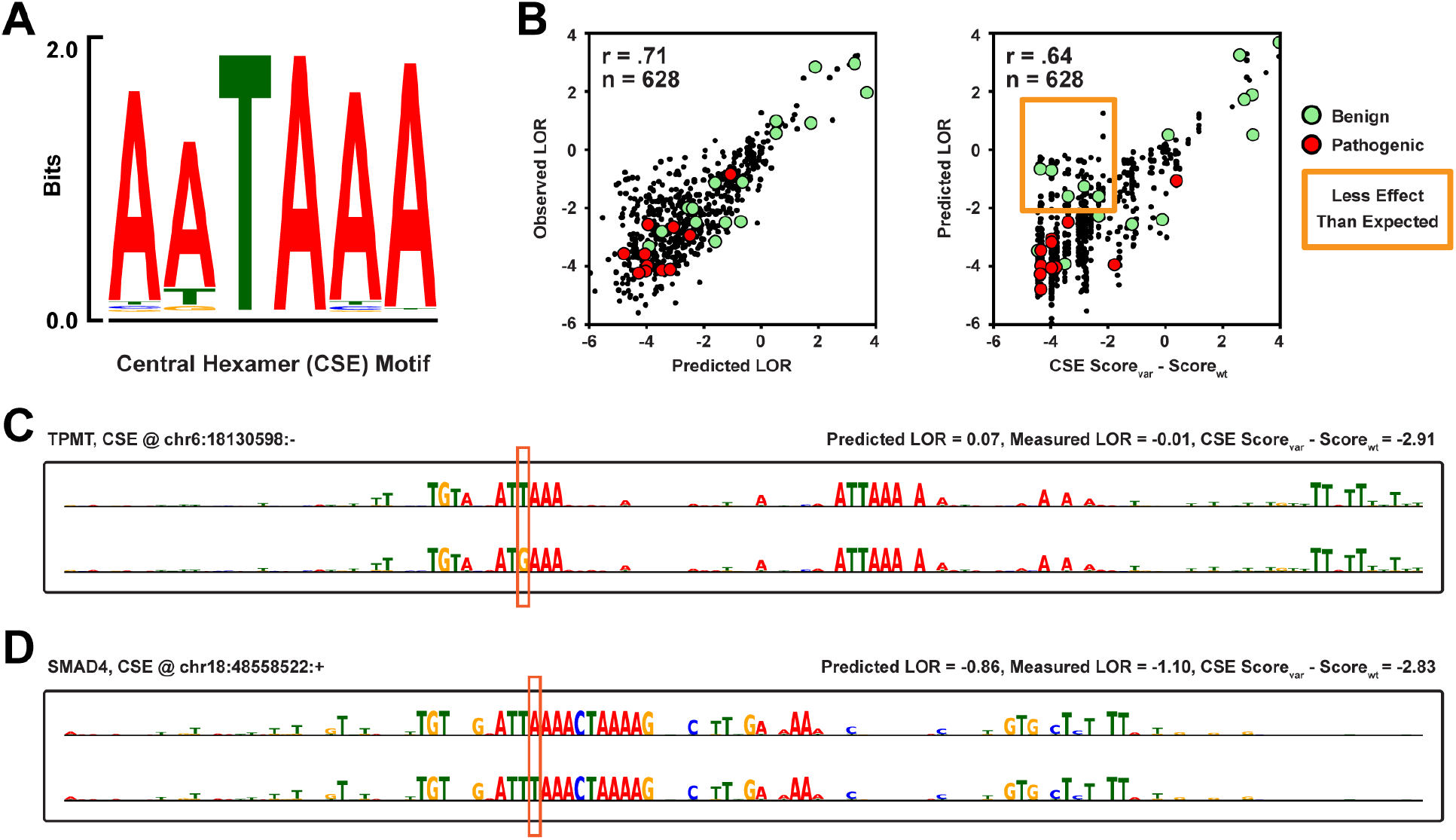
**A** Position weight matrix (PWM) of the CSE motif, as measured in the MPRA of Bogard et al. [36]. **B** Predicted vs measured log odds ratio of CSE mutations from the MPRA (n = 628). Right: Log odds ratio predicted by APARENT2 vs the effect sizes predicted by a linear hexamer model trained on the same data. **C** Interpretation of a functionally silent CSE mutation in the TPMT gene. **D** Interpretation of variant with dampened effect size in the SMAD4 gene.

To identify CSE variants with predicted effect sizes lower than expected, we compared APARENT2’s predictions to a linear CSE hexamer regression model trained on the same data (Figure 4B, right) [36]. While the two models generally agree (spearman r = 0.64), we find multiple mutations with log odds ratios *< −*2 as predicted by APARENT2 but with log odds ratios *> −*2 as predicted by the hexamer model. All variants that occur in ClinVar within this group are labeled benign. Using our mask-based interpretation method, we dissected the origin of this discrepancy. First, we find a group of completely silent mutations and these PASs all contain redundant CSE hexamers (Figure 4C, Supplementary Figure S4A). Importantly, the interpretations show that besides the extra CSE motifs, it is crucial that auxiliary elements (e.g. CFIm-binding TGTA motifs or downstream T-rich elements) are well-positioned with respect to the new CSE. Second, we find another group of variants with dampened effect sizes when mutating the canonical CSE into a weaker form (Figure 4D, Supplementary Figure S4B). Rather than redundant CSE motifs, these PASs contain many well-positioned auxiliary motifs (CFIm- and CstF-binding motifs and T-rich elements) which dampen the loss of the canonical CSE hexamer. This hypothesis agrees with earlier work suggesting that weak CSEs are efficient polyadenylation elements when found in a strong sequence context [66, 67].

### Disruptive Polyadenylation Variants are Selected Against in the Human Population

We next sought to understand the connection between the functional impact of genetic variation on polyadenylation and human health. Using APARENT2 we performed full in-silico saturation mutagenesis of every annotated PAS in PolyA DB V3 [51, 52] and imputed the effect size (odds ratio) of every possible SNV (n *>*43.8 million). For each PAS, we calculated the average wildtype isoform usage across all tissues in PolyA DB. We then re-calculated the isoform usage in the presence of each mutation by using the APARENT2 prediction to scale the isoform odds. Given these two quantities we estimated the change in isoform proportion (Δuse) due to each variant. When cross-referencing our predictions against the *>*2, 8 million PAS SNVs curated from the *>*71, 000 genomes sequenced in gnomAD v3 [48] (Figure 5A), we found that disruptive loss-of-function variants (resulting in downregulated pA) are depleted in common variants (AF *>*0.1%) compared to singletons (wilcoxon *p* = 2.1*×*10^*−*76^; Figure 5B, Supplementary Figure S5A). Disruptive loss-of-function variants (Δuse *< −*0.15) occur *∼*2.5-fold less frequently among common variants (AF *>*10%) than singletons and they occur *∼*1.4-fold more frequently in unobserved variants (AF = 0%) compared to singletons (Figure 5C). These results suggest a negative selection pressure on disruptive variants in human polyadenylation signals.

**Figure 5:**
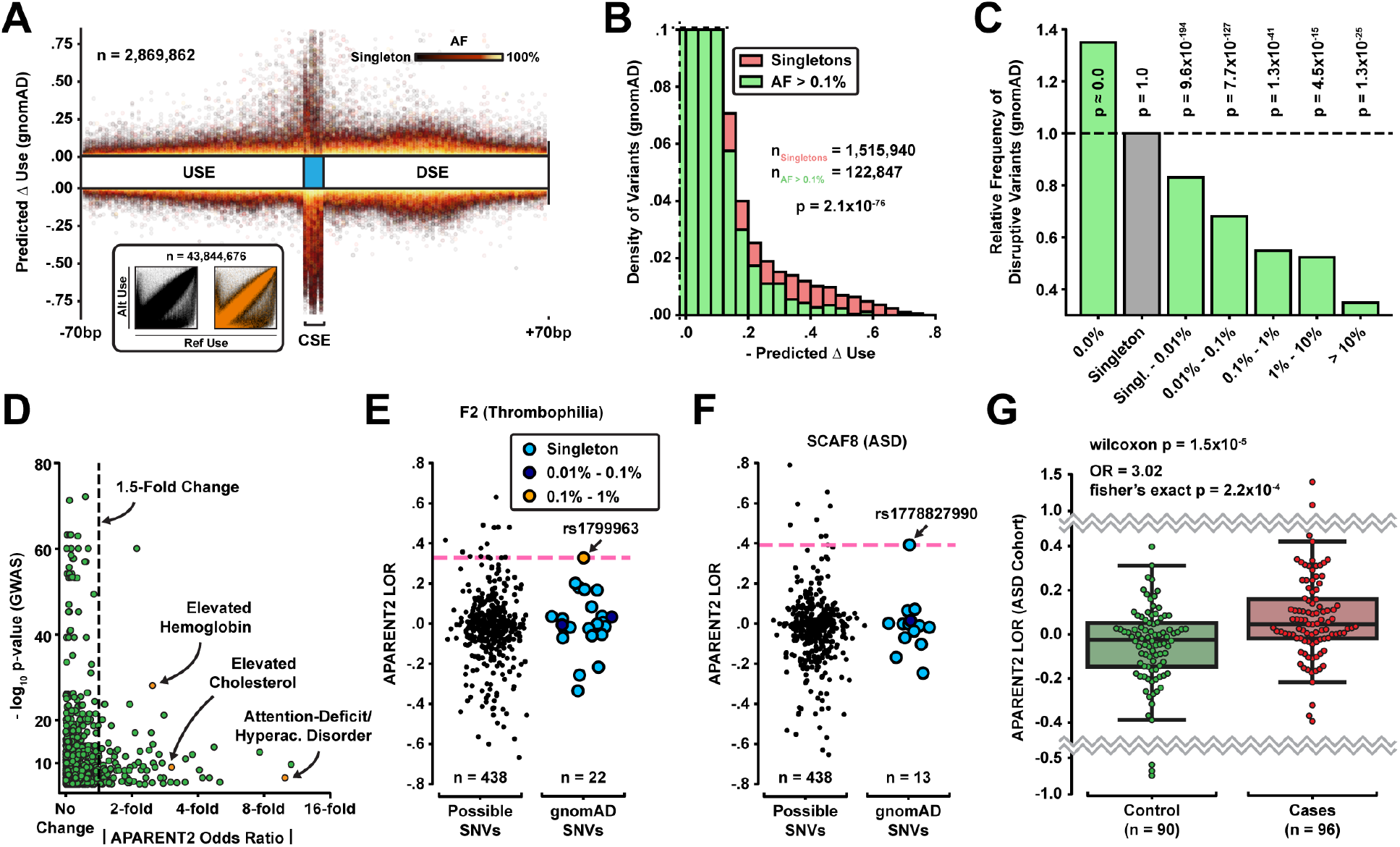
**A** Relative position of mutation vs predicted Δ isoform abundance for all PAS variants in gnomAD (n = 2.8 million). Color intensity represents allele frequency. Inset: Reference vs alternate isoform abundance for all 43.8 million potential PAS SNVs. **B** Distribution of predicted Δ isoform abundance for common gnomAD variants (AF *>*0.1%; green) and singletons (red). **C** Relative frequency of disruptive variants (Δuse *< −*0.15) with respect to singleton variants. Shown above each bar are the corresponding wilcoxon p-values. **D** Absolute predicted isoform fold change vs p-value (-log_10_) of GWAS Catalog SNPs (n = 1, 233). **E** Distribution of predicted log odds ratios for the F2 PAS. **F** Distribution of predicted log odds ratios for the SCAF8 PAS. **G** Distribution of predicted log odds ratios among ASD cases and controls from a WGS study [49].

### Gain-of-Function Mutations in the 3’-End are Associated with Clinical Conditions

Most polyadenylation variants that have previously been associated with disorders are highly disruptive and rare CSE mutations [16, 17, 18]. However, while we found in the previous section that highly disruptive loss-of-function variants are generally selected against, they also frequently occur as common variants. This suggests that we cannot use the inferred effect on polyadenylation alone as a predictor for variant pathogenicity. For example, when we intersected our transcriptome-wide PAS variant predictions against the GWAS catalog (n = 1, 233) we observed common SNPs with both large (*>*1.5-fold) and small (*<*1.5-fold) predicted effect sizes that are strongly associated with phenotypes such as elevated cholesterol and Attention-deficit / Hyperactivity Disorder (Figure 5D) [68]. Similarly, the weak gain-of-function mutation 97G*>*A in the PAS of the F2 gene (a variant that increases pA efficiency *<* 1.5-fold) is responsible for Thrombophilia, a highly penetrant hypercoagulable condition in humans [19]. Clearly, the downstream consequence of disrupted polyadenylation depends on the regions of the 3’ UTR that are affected by the APA isoforms, not to mention the gene itself. However, we can assume that a mutation is likely not deleterious if it occurs in a PAS with common variants that have even larger effect sizes. Thus, we can eliminate PAS mutations and classify them as likely benign when they co-occur with putative functional common variants in gnomAD with high impact on polyadenylation. For example, the known pathogenic F2.*97G*>*A mutation would not be eliminated, since it is the variant with largest predicted odds ratio of all observed variants in gnomAD (Figure 5*E*).

Using the stratification process above, we investigated the link between misregulated polyadenylation and Autism Spectrum Disorder (ASD), a relationship which has been suggested before but mainly at the trans-regulatory level and less in terms of cis-regulatory variation in the 3’ UTR [69, 70, 71, 72]. Figure 5F displays an example rare variant (rs1778827900) associated with ASD [73]. The suspected variant has a considerably higher (positive) effect size than any of the observed variants in gnomAD. Hypothesizing that gain-of-function mutations may be linked to ASD, we ran APARENT2 on whole-genome sequencing (WGS) data from 1, 902 families [49] and found that variants overlapping PASs in cases are enriched for gain-of-function compared to controls (wilcoxon *p* = 0.049, *n*_cases_ = 297, *n*_controls_ = 296). When removing variants that co-occur with higher-impact common SNPs in gnomAD (AF *>*0.01%), the significance increased (wilcoxon *p* = 2.1*×*10^*−*4^), and when also removing variants that occur in PASs with a protective downstream PAS within 200nt, the significance increased further (wilcoxon *p* = 1.5*×*10^*−*5^; see Methods for filtering procedure) (Figure 5G, Supplementary Figure S5B). We observed a 3.02-fold enrichment of gain-of-function mutations in cases (fisher’s *p* = 2.2*×*10^*−*4^). As additional validation, the predicted effect sizes of variants from the control set were indistinguishable from variants in gnomAD after applying the same filtering (wilcoxon *p* = 0.341) while case variants were significantly different (wilcoxon *p* = 2.7*×*10^*−*5^). Finally, we found an enrichment among PAS case variants of the gene ontology terms ‘regulation of primary metabolic process’ (FDR = 6.79*×*10^*−*02^) and ‘protein binding’ (FDR = 3.48*×*10^*−*04^) [74]. We found no significant enrichment among controls.

When replicating our analysis against the smaller WGS study of 200 families from Yuen et al. [73], we again observed an enrichment of gain-of-function mutations in cases relative to the controls from An et al. [49], but the results were only significant with less stringent filtering criteria (wilcoxon *p* = 0.039; Supplementary Figure S5C-D; see Methods). The predicted effect sizes of case variants were not significantly different from gnomAD variants (wilcoxon *p* = 0.127), but the trend was similar to that of the larger cohort data from An et al. [49] so this is likely due to insufficient sample size of the smaller WGS study. Even in this smaller cohort, we can use APARENT2 to functionally interpret variants with high predicted effect sizes. For example the variant highlighted in Figure 5F (rs1778827900) is found to be a gain-of-CstF mutation with super-additive interactions to neighboring T-rich elements (Supplementary Figure S5E).

## Discussion

In this paper, we developed an improved human polyadenylation variant prediction model, APARENT2, based on deep residual neural networks. We systematically compared this model to other sequence-predictive APA models, including the original APARENT network, on the task of predicting functionally disruptive variants from MPRA data and human APA QTLs. We found that APARENT2 was considerably better at variant effect size estimation compared to other models, in particular for cryptic variants outside of the CSE. We further trained tissue-specific residual models for testis, ovary, B-cell lymphocytes and brain and used these to improve variant prediction in human tissues. By combining rich modeling with mask-based attribution, we extracted complex cis-regulatory rules and elucidated cooperativity among core polyadenylation signal motifs. For example, we found super-additive interactions between the CFIm-binding motif TGTA and downstream AU-rich elements. Conversely, we identified protective buffering effects of redundant and well-positioned core hexamers that can ‘take over’ in case the original CSE is disrupted by mutations.

An intriguing finding of our work is that the same PAS scoring function accurately predicts relative isoform abundance in multi-PAS genes and absolute transcript levels in genes containing a single PAS. These results are consistent with a simple model of polyadenylation where a PAS emerging during transcription is used with an independent probability that is determined entirely by the sequence of that signal. If an additional PAS occurs in the emerging transcript, its usage is again determined independently by the sequence. Moreover, 3’-end processing via cleavage and polyadenylation is in competition with other processes such as RNA degradation and transcriptional feedback that reduce mature mRNA levels.

We applied APARENT2 to make functional predictions on 44M PAS variants in the human genome, orders of magnitude more than would currently be possible even in a high-throughput reporter assay. Moreover, unlike statistical methods such as aQTL analysis, functional predictions can be made even for variants that have not yet been observed, but may well occur, in the human population. Finally, we combined APARENT2’s variant predictions with additional evidence from a large variation database (gnomAD). This allowed us to enrich our predictions by disregarding mutations that co-occur in PASs with common high-impact variants, as these PASs are likely not important for function. Using this approach, we found a *∼*3-fold enrichment of gain-of-function variants (leading to more efficient pA) in individuals with Autism Spectrum Disorder.

It is important to note that we cannot definitively classify mutations in PASs with high predicted effect size as causative of Autism; both loss- and gain-of-function variants occur frequently in controls, suggesting many gain-of-function variants in cases are likely benign. However, the fact that there is a significant over-representation of gain-of-function mutations in cases suggests that *some* of those variants contribute to Autism. Using our predictions, we can propose outlier variants for experimental validation, for example by quantifying isoforms in cases from RNA-seq data or with functional screens in mouse models. These results signify the importance of having a functional model; the number of mutations occurring in PASs were almost identical between cases and controls and we only discovered the signal in APARENT2’s predictions.

## Methods

### Neural Network Architecture

APARENT2 is based on residual blocks of dilated convolutions [50] and is architecturally similar to the SpliceAI model [27]. Let *𝒫* be the APARENT2 model. As input, *𝒫* receives a one-hot coded sequence ***x*** *∈* {0, 1}^205*×*4^, which represents the proximal PAS, and a one-hot coded variable ***l*** *∈* {0, 1}^13^ which indicates the source 3’ UTR sub-library from the MPRA training data [36]. Internally, *𝒫* consists of 7 **Residual Groups**, and each residual group is made up of 4 **Residual Blocks**. A residual block (Supplementary Figure S1A) consists of two batch-normalized, ReLU-activated one-dimensional convolutional layers with a specific filter dilation rate. Each block also has a skip connection, which mathematically performs an unweighted element-wise addition. Each residual group consists of residual blocks of the same dilation rate. For this particular network, the 7 residual groups use the following sequence of dilation rates: 1, 2, 4, 8, 4, 2, 1. Between each residual group, there is an extra skip connection to the final output layer. Only ***x*** is passed through the series of residual blocks, producing in the end a single-channel vector of non-normalized cleavage scores ***s***(***x***) *∈* ℝ^206^ (Note that ***s*** has one position more than ***x***; this extra position represents the total isoform score of the distal signal). The library indicator variable ***l*** is multiplied with a position-specific weight matrix ***W*** *∈* ℝ^206*×*13^ and linearly combined with ***s***(***x***), producing new scores 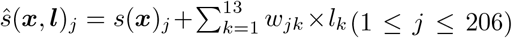 which have effectively been scaled with a library-specific intercept. Finally, *𝒫* produces a normalized 206-way cleavage distribution ***ŷ*** *∈* [0, 1]^206^ by applying the softmax transform (Equation 1).

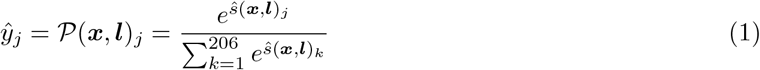

All residual blocks in APARENT2 have 32 channels and all convolution filters are 3 positions wide. Note that there is no explicit sigmoid output representing the total proximal isoform proportion. Rather, the proximal isoform proportion is computed as the sum of cleavage probability mass 7–57 nt downstream of the start of the proximal CSE (which is located at position 70): 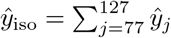. However, for some variant prediction tasks the proximal isoform is defined as ‘any cleavage that is not distal’ (i.e. the data processing of those datasets considered cleavage from nearby competing cryptic PASs as ‘proximal’). In that case, we define the predicted proximal isoform proportion as 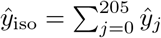 (and *ŷ*_206_ is the distal proportion).

### MPRA Training Data

The MPRA dataset from Bogard et al. [36] was re-processed to make the training data more uniform. First, the original dataset consisted of 185 nt long sequences, starting 50 nt upstream of the proximal CSE. However, for some of the sub-libraries (the MPRA consisted of 13 sub-libraries with different 3’ UTR contexts), an additional random barcode was located from 70 to 50 nt upstream of the CSE. In the re-processed version of the data, we included 20 nt of additional sequence upstream of the CSE to capture these barcodes.

Second, for some of the sub-libraries, the exact cleavage distributions were not estimated from the RNA-Seq data. Instead, these sub-libraries only included total proximal-to-distal isoform proportions. We re-mapped the RNA-Seq reads to these sub-libraries and augmented the data with the missing cleavage distributions.

Finally, the original models were only trained on about 2.4 million of the degenerate (randomized) MPRA data (3 of 12 sub-libraries of the random MPRA were held out for independent testing), and it was not trained on any of the assayed human APA sites from the designed MPRA. Here, we trained the network on data from all of the degenerate sub-libraries, resulting in 3.3 million training sequences and 80, 000 sequences for validation and testing each. We also included human intronic PAS sequences from the designed MPRA (which had been measured in a 3’ UTR reporter), adding approximately 10, 000 additional high-quality measurements to the training data. To keep the variant prediction results unbiased, we did **not** train the network on any of the human 3’ UTR sequences from the variant MPRA. Note that, as in the original paper, MPRA sequences with *>*75% adenine bases in a 12-20bp region were removed to minimize internal priming artifacts [36]. Hence, the resulting trained model can not be used on sequences with long adenine stretches.

### Cleavage and Isoform Cost Function

Given the training data 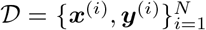, where ***x*** *∈* {0, 1}^205*×*4^ is a one-hot coded representation of the proximal (degenerate) polyadenylation signal and ***y*** *∈* [0, 1]^206^ is the measured 3’ cleavage distribution, we trained APARENT2 to minimize the hybrid cost function given in Equation 2.

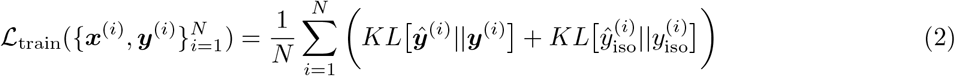

Here, *KL* [***ŷ***^(*i*)^||***y***^(*i*)^] is the KL-divergence between the predicted cleavage distribution ***ŷ***^(*i*)^ = *𝒫* (***x***^(*i*)^, ***l***^(*i*)^) and measured distribution ***y***^(*i*)^ (defined in Equation 3). 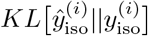 is an extra consistency term used to fit the sum of a subset of softmax outputs 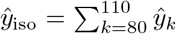to the total observed proximal isoform proportion *y*_*k*_ (defined in Equation 4). Note that the last position in the target vectors 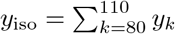 corresponds to the total isoform proportion (cumulative cleavage) of the distal (non-degenerate) PAS.

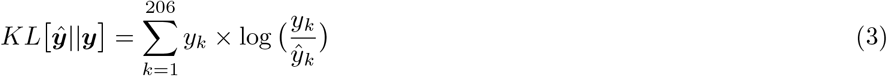

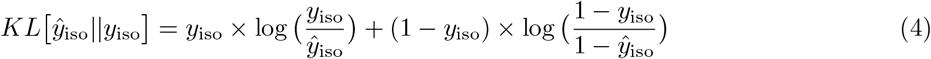

### Model Training

We trained APARENT2 for 5 epochs with mini-batch SGD using the Adam optimizer in Keras [75, 76] with default parameters and batch size = 64. During training, we randomly shift both the input sequence ***x***^(*i*)^ and target cleavage distribution ***y***^(*i*)^ by at most 15 nt in either direction (Supplementary Figure S1B). As such, the CSE position (which all sequences are initially aligned against) varies during training, forcing the network to learn to displace the cleavage distribution according to the location of the CSE. This helped the network give better predictions to locally competing CSE hexamers in the nearby USE or DSE regions.

### Web Tool

We developed a web tool for running in-silico saturation mutagenesis across human polyadenylation signals from the PolyA DB V3 data [51, 52] (Supplementary Figure S6). The application loads a graph tool based on D3.JS [77], where predicted cleavage distributions can be explored interactively. *Note:* This web application has been online since 2019, but has been relying on the original APARENT model for predictions.

### Endogenous Datasets

We collected three different sets of human 3’-end sequencing data in order to benchmark the APA models at the tasks of predicting pairwise and multi-PAS isoform proportions. We first downloaded the tissue-pooled version of APADB [56] from http://tools.genxpro.net:9000/apadb/download/track/hg19.apadb_v2_final.bed/ (dataset # 1). We then downloaded the RNA-seq counts of Lianoglou et al. [41] from https://cbio.mskcc.org/leslielab/ApA/atlas/ and mapped the read positions to the annotated PASs in APADB. From the mapped cleavage position counts, we estimated the total read count 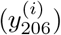 that support APA isoform *i* of gene *k* aggregated over all tissues (dataset # 2). Similarly, the aggregated isoform counts from Derti et al. [13] were downloaded from GEO (accession GSE30198) and mapped to the PAS sequences from APADB (dataset # 3).

For the task of pairwise APA isoform prediction, we collected pairs of adjacent PASs with a total read count *≥*500. The two sites had to be at least 100bp apart and at most 4, 000bp apart. Finally, sequences with more than 7 consecutive adenine bases were removed to minimize the risk of internal priming. For the multi-PAS prediction task, we kept genes with at least 2 annotated PASs in APADB and at most 10 PASs. We removed genes with less than 10 total counts, or with PASs separated by less than 50*bp* or more than 40, 000bp. Genes with PASs that contain more than 13 consecutive adenines were removed.

### Variant Datasets

We benchmarked the APA models on two 3’ UTR MPRAs and two native transcriptomic 3’ aQTL datasets. The specific data filters and measurements of each dataset are described below.

### Isoform MPRA

(Bogard et al. [36]) This APA variant MPRA contains SNVs near PASs of disease-implicated 3’ UTRs from ClinVar, HGMD or ACMG genes [38, 39, 40]. We filtered the data to include only variants where the wildtype and variant sequences each had a mean unique UMI read count *>* 200 from at least 5 barcoded replicates. This resulted in a total of 12, 350 retained variants. We estimated the log odds ratio (log fold change) LOR(*y*_wt_, *y*_var_) of each variant’s proximal isoform abundance *y*_var_ with respect to the wildtype abundance *y*_wt_ (Equation 5). These isoform abundances were calculated by summing all cleavage probabilities mapping to cut sites +0 to +50 nt downstream of the CSE.

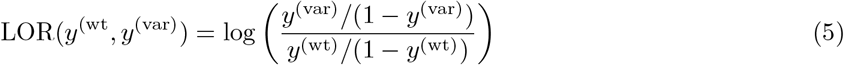

In one of the benchmarks, we compared the model performances of classifying disruptive APA variants. A variant was deemed ‘disruptive’ if the absolute value of its isoform odds ratio with respect to the wildtype abundance was larger than 2: Disruptive = 1 if 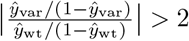 else 0.

The dataset was downloaded from: https://github.com/johli/aparent.

### Expression MPRA

(Slutskin et al. [35]) This 3’ UTR MPRA measured the RNA/DNA fold changes of viral PAS variants. We filtered the data to include only sequences that were in the test set of the PolyApredictor model. We also removed sequences that contained either a stretch of at least 10 consecutive A’s, or sequences containing the subsequence ‘AGA’ at position 41, as the measurements of these sequences seemed to be influenced by artifacts. Finally, we only considered the subset of sequences that were part of the scanning mutagenesis experiments, resulting in a total of 442 variants. For these sequences, we matched the wildtype and variant PASs in order to calculate the RNA/DNA fold change ratio *FCR*(*u*_wt_, *u*_var_) = *u*_var_ *−u*_wt_ due to each variant. Here *u* is the logarithm of the RNA/DNA fold change of a particular sequence.

The model PolyApredictor predicts the log fold changes *û* directly. For all other models, we approximate *û* with the predicted isoform log odds log (*ŷ/*(1*−ŷ*)). Furthermore, since both PolyApredictor and APARENT2 supports cleavage predictions at base-pair resolution, we also compared them on their ability to infer the RNA/DNA fold change ratios *FCR*_*j*_ of each variant across every wildtype cleavage position *j*.

The dataset was downloaded from: https://github.com/segallab/PolyApredictors.

### GTEx 3’ aQTLs

[20] The GTEx v7 3’ aQTL data was downloaded and mapped to the PolyA DB V3 annotation [51, 52]. The data was further filtered to only include Lead SNPs occurring within 50nt of the most likely CSE of the annotated PAS. This resulted in 366 SNPs with measured 3’ aQTL effect sizes among 44 GTEx tissue types. To predict effect sizes, we first used each model to infer the SNP log odds ratio LOR(*ŷ*^(wt)^, *ŷ*^(var)^). Next, given the observed Polyadenylation Distal Usage Index *y*_PDUI_ *∈* [0, 1] of a particular PAS averaged across all GTEx tissues and samples, we inferred the SNP effect size Δ*y*_PDUI_ by scaling the measured PDUI with the predicted variant odds ratio (Equation 6).

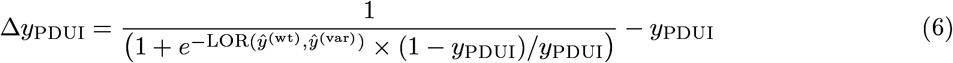

The dataset was downloaded from: https://doi.org/10.7303/syn22236281.

*Note:* In Supplementary Figure S3A, we use APARENT2 to predict variant log odds ratios for all cis-acting GTEx v8 SNPs overlapping PASs from Cui et al. [78], but we did not use this data in the benchmark analysis since only the p-values are publicly available, not the effect sizes nor lead SNP classifications.

### HapMap Yoruba Lymphoblastoid 3’ aQTLs

[21] The 3’ aQTLs were mapped against the PolyA DB V3 annotation [51, 52] and narrowed to the subset of SNPs occurring within 50nt of the most likely CSE of each annotated PAS. This resulted in 58 variants measured among 52 HapMap Yoruba human lymphoblastoid cell lines. We used the effect sizes estimated from nuclear mRNA only. The predicted log odds ratio LOR(*ŷ*^(wt)^, *ŷ*^(var)^) of each model was directly compared to the 3’ aQTL effect sizes.

The raw data and annotations were available at GEO under accession GSE138197. The processed data, including the estimated 3’ aQTL effect sizes, were provided to us by the authors.

### Variant Prediction Models

Following is a list of the APA models that were included in the variant prediction benchmark, with a detailed description of how each model was used and where each model was downloaded from.

#### APARENT2

(This paper) The model takes as input a 205 nt one-hot coded sequence ***x*** *∈* {0, 1}^205*×*4^ and a MPRA sub-library indicator ***l*** *∈* ℝ^13^. The model predicts a 3’ cleavage distribution ***ŷ*** *∈* ℝ^206^ (*ŷ*_206_ corresponds to total isoform cleavage). When using the model for variant prediction, we set *l*_11_ = 1 (the human intronic PAS sub-library intercept). The variant log odds ratio LOR(*ŷ*^(wt)^, *ŷ*^(var)^) is calculated from a subset of the cleavage outputs. For the MPRA of Bogard et al. [36] and the 3’ aQTLs of Mittleman et al. [21], we define 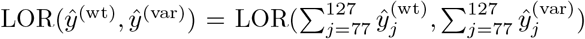. For the MPRA of Slutskin et al. [35] and the GTEx 3’ aQTLs [20], we define 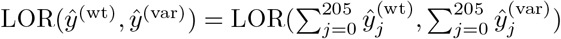.

#### APARENT

(Bogard et al. [36]) The original APARENT model, which takes as input a 185 nt one-hot coded sequence ***x*** *∈* {0, 1}^185*×*4^, a MPRA sub-library indicator ***l*** *∈* ℝ^13^ and a binary variable *d ∈* {0, 1} which indicates whether there is a far-away distal PAS in the MPRA sub-library. The model produces two outputs, a total proximal isoform proportion *ŷ*_iso_ *∈* ℝ, and 3’ cleavage distribution ***ŷ*** *∈* ℝ^186^ (*ŷ*_186_ corresponds to total isoform cleavage). When using the model for variant prediction, we set *l*_4_ = 1 and *d* = 1. The variant log odds ratio LOR(*ŷ*^(wt)^, *ŷ*^(var)^) is calculated as the average of the isoform- and cleavage outputs: 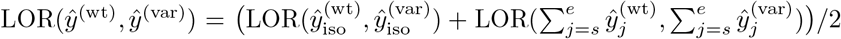, where *s* = 57 and *e* = 107 for the MPRA of Bogard et al. [36] and the 3’ aQTLs of [21], and *s* = 0 and *e* = 205 otherwise.

The trained model was downloaded from: https://github.com/johli/aparent/tree/master/savedmodels.

#### DeeReCT-APA

(Li et al. [34]) An LSTM-based model trained on mouse 3’-sequencing data. The model takes as input a tensor ***x*** *∈* {0, 1}^*P ×*455*×*4^, where ***x***_*p*_ *∈* {0, 1}^455*×*4^ denotes the *p*:th PAS in a given 3’ UTR. When using the model for SNV prediction, we only pass two input PASs (*P* = 2) – the sequence of the PAS containing the mutation and a fixed distal PAS that we never change. The distal PAS was chosen as a strong sequence from the training data. We could not use the distal PAS from the variant MPRA, since the model’s input window was larger than the plasmid reporter 3’ UTR. By passing either the wildtype- or variant sequence as the proximal PAS, the model returns the predicted wildtype- and variant isoform proportions *ŷ*^(wt)^ and *ŷ*^(var)^. Given these predictions, we calculate LOR(*ŷ*^(wt)^, *ŷ*^(var)^).

The model was re-trained using the code from: https://github.com/lzx325/DeeReCT-APA-repo.

#### DeepPASTA (Rel Iso)

(Arefeen et al. [33]) An ensemble of CNNs trained on human 3’ sequencing data [13]. We used the ‘Tissue-specific, relatively dominant’ models. These tissue-specific model instances take as input two 200 nt one-hot coded sequences ***x***^(*p*)^, ***x***^(*d*)^ *∈* {0, 1}^200*×*4^ (proximal and distal PAS) and one-hot coded representations ***s***^(*p*)^, ***s***^(*d*)^{0, 1}^200*×*7^ of their most probable secondary structures. When using the models for SNV prediction, we use a fixed distal PAS that we never change. By passing either the wildtype- or variant sequence as the proximal PAS, the model returns the predicted wildtype- and variant isoform proportions *ŷ*^(wt)^ and *ŷ*^(var)^. Given these predictions, we calculate LOR(*ŷ*^(wt)^, *ŷ*^(var)^). This is repeated for each tissue-specific model and the average LOR is used as the final prediction.

The trained models were downloaded from: https://www.cs.ucr.edu/ãaref001/DeepPASTAsite.html.

#### DeepPASTA (Site Pred)

(Arefeen et al. [33]) This CNN ensemble only takes a single one-hot coded sequence ***x*** *∈* {0, 1}^200*×*4^, and one-hot coded representations ***s***^(1)^, ***s***^(2)^, ***s***^(3)^{0, 1}^200*×*7^ of the three most probable secondary structures, as input. The model predicts the likelihood of ***x*** being a PAS. By passing either the wildtype- or variant sequence as ***x***, the model returns the predicted wildtype- and variant PAS probabilities *ŷ*^(wt)^ and *ŷ*^(var)^. Given these predictions, we calculate LOR(*ŷ*^(wt)^, *ŷ*^(var)^).

The trained model was downloaded from: https://www.cs.ucr.edu/ãaref001/DeepPASTAsite.html.

#### PolyApredictor

(Slutskin et al. [35]) An RNA/DNA expression level CNN and a 3’-cleavage CNN (as two separate networks) trained on a plasmid reporter MPRA of 3’ UTRs (assayed in K562 cells). The models each take a one-hot coded sequence ***x*** *∈* {0, 1}^250^ as input, which represents the 3’ UTR, and predicts either the total 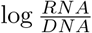level *ŷ ∈* ℝ or the per-nucleotide 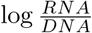 levels ***ŷ*** *∈* ℝ^250^ across all potential cleavage positions of the 3’ UTR. To use these models for variant prediction, we pass either the wildtype- or variant sequence as ***x*** and calculate the predicted LOR as: LOR(*ŷ*^(wt)^, *ŷ*^(var)^) = *ŷ*^(var)^ *− ŷ*^(wt)^.

The trained models were downloaded from: https://github.com/segallab/PolyApredictors.

### Motif Discovery

To generate a representative selection of RNA binding protein motifs within human polyadenylation signals, we used APARENT2 to predict the isoform logit log *ŷ*_iso_*/*(1*−ŷ*_iso_) for 20, 000 randomly sampled 3’ UTR PASs from PolyADB V3 (where 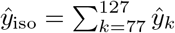). The PASs were restricted from having more than 7 consecutive adenine bases, the signals had to be alternatively used in tissue-pooled measurements and the CSE had to have a hamming distance of at most 2nt from the consensus AATAAA motif. We used DeepSHAP [79] to obtain attribution scores for each PAS (64 reference patterns), which were clustered into sequence logos using TF-MoDISco [54] (sliding window = 8, flank size = 5, max seqlets = 40, 000, FDR = 0.05, # mismatches = 0). The TF-MoDISco software was installed from https://github.com/kundajelab/tfmodisco/.

### Pairwise and Multi-PAS Modeling

All APA models were benchmarked on the pairwise APA prediction task using the three endogenous data sources from Müller et al. [56], Derti et al. [13] and Lianoglou et al. [41]. For each dataset, we estimated the true isoform logit of every pair of APA sites as logit_endogenous_ = logit (*c*_p_ + *c*_pseudo_)*/*(*c*_p_ + *c*_d_ + *c*_pseudo_), where *c*_p_ and *c*_d_ are the proximal and distal isoform counts and *c*_pseudo_ = 0.5 is a pseudo count. We then used each APA model to predict logit scores logit_p_ and logit_d_. These scores were used together with the log-distance *d* between the sites to regress the estimated logits from the endogenous data (Equation 7):

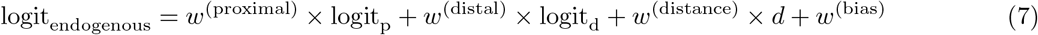

For the multi-PAS task, we estimated the distal isoform proportion of each gene as 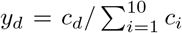. We then feed each APA model the 10 input PAS sequences of the gene (with zero-padding if the gene has less than 10 PASs). Each model returns 10 predicted logit scores logit_*i*_, which are used in a masked softmax regression model to predict the distal isoform proportion of the endogenous data (Equation 8):

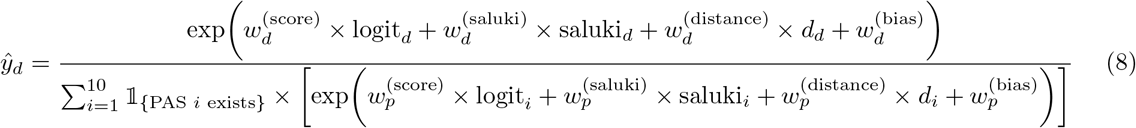

The variable *d*_*i*_ denotes the cumulative log distance between PAS *i* and the distal-most PAS in the equation above. The variable saluki_*i*_ denotes the mean 50-fold ensemble prediction of the Saluki model for isoform *i* [55]. Saluki was downloaded from https://zenodo.org/record/6326409. Each 3’ UTR isoform was extracted from the GENCODE v19 annotation starting from the last defined stop codon of each gene and ended at the median cleavege site of the PAS [80]. A constant 5’ UTR and ORF, taken from https://github.com/vagarwal87/saluki_paper, were used for all 3’ UTRs. Note that Saluki inputs were only used for the model named ‘APARENT2+Saluki’ in the benchmark of Figure 1G. The parameters of the softmax regression model were fit using LM-BFGS. In one of the tests in Figure 1G, the APARENT2 logits, Saluki scores, PAS log distances and a 0*/*1-vector indicating the distal PAS was used to fit a single-layer LSTM model with 16 hidden units instead of the softmax regression model of Equation 8. This model was trained in Keras with 20-fold cross-validation [75].

### Tissue-Specific Modeling of Native APA

Here we again used the tissue-specific 3’ RNA-seq data from Lianoglou et al. [41], but rather than aggregating isoform counts over tissues, we now keep track of the total read count 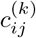supporting APA isoform *i* in tissue *j* of gene *k*. We removed genes with more than 10 APA isoforms. We estimated isoform proportions by normalizing the isoform counts by the total count across each gene: 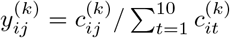. We then created separately filtered copies of the data for pairs of tissues, where one tissue was HEK293 and the other tissue was either testis, ovary, BLCL or brain. Genes with less than 10 total supporting reads in any tissue were removed from each dataset. This resulted in 4, 453 (HEK293 – Testis) genes, 4, 495 (HEK293 – Ovary) genes, 4, 366 (HEK293 – BLCL) genes and 4, 715 (HEK293 – Brain) genes.

Using these data, we trained 4 individual tissue-specific models that learn the difference in isoform proportion 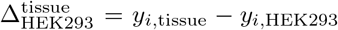 between the target tissue type and HEK293. The model works as follows: Given the 10 input PAS sequences ***x*** *∈* {0, 1}^10*×*205*×*4^ of a given gene (with appropriate zero-padding), we execute APARENT2 on each PAS to obtain baseline cleavage predictions ***ŷ*** *∈* [0, 1]^10*×*206^. We compute the baseline isoform logit for PAS *i* as 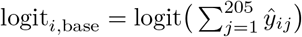. We also feed ***x*** as input to a trainable CNN that predicts tissue-specific scores ***ŝ*** *∈* ℝ^10*×*2^. The CNN weights are shared across all 10 PASs. Internally, the tissue-CNN consists of 2 convolutional layers (16 filters, 8 positions wide) and global average pooling. Finally, we linearly combine logit_*i*,base_ and *ŝ*_*i*,tissue_ with the log distance *d*_*i*_ between PAS *i* and the distal-most PAS and apply masked softmax to predict tissue-specific isoform proportions *ŷ*_*i*,tissue_ (Equation 9).

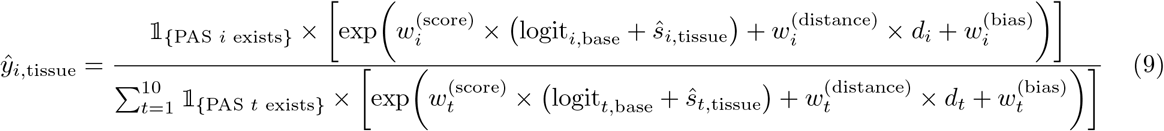

For the first 25 epochs, we froze the tissue-CNN weights and forced *ŝ*_*i*,tissue_ to be 0. We only optimized the PAS-specific regression weights ***w*** *∈* ℝ^10*×*3^ from Equation 9. We minimized the mean (masked) KL divergence between predicted and observed tissue-specific isoform proportions across all *K* genes (Equation 10). Consequently, the regression weights ***w*** will learn to combine the baseline APARENT2 scores and the log distances to infer native isoform proportions.

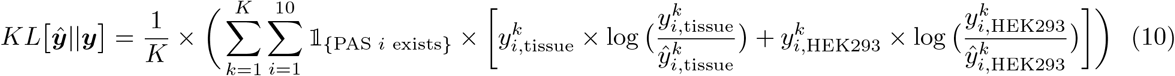

After the weights ***w*** converge, we un-froze the tissue-CNN weights and optimized ***w*** jointly with the CNN-predicted scores *ŝ*_*i*,tissue_ for 25 additional epochs. We noticed that if we keep minimizing the KL-divergence loss of Equation 10, the CNN would disregard learning about tissue-specific differences (which is a relatively small source of variation for APA) in favor of learning to better predict the mean proportion across both tissues. We thus switched to a (masked) margin loss which penalized the model based on the observed and predicted tissue-specific differences 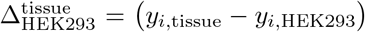 and 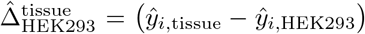 rather than absolute proportions (Equation 11).

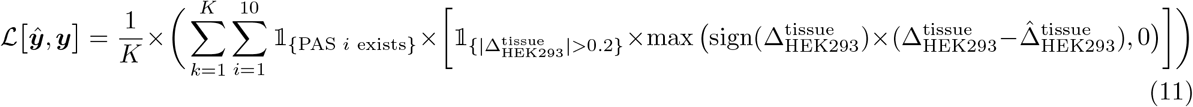

For each tissue-specific model (testis, ovary, BLCL, brain), we learned an ensemble of 10 independently trained CNNs. We stopped when the validation error on a held-out test set of 500 genes started to increase.

### Tissue-Specific aQTL Effect Size Prediction

We used the testis-, ovary-, BLCL- and brain-specific APA models to scale the effect size predictions made by APARENT2 on the GTEx aQTLs [20]. We used the linear model proposed by Cheng et al. [60] for combining baseline variant predictions with a tissue-specific scaling factor. Specifically, the logit of the tissue-specific distal isoform usage (PDUI) for the variant sequence is assumed to follow the relationship of Equation 12.

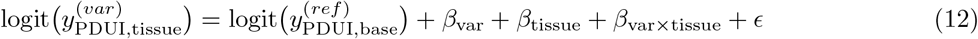

Here 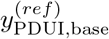 corresponds to the mean PDUI measured across all tissues and samples. If we set

- *β*_var_ = LOR(*ŷ*^(wt)^, *ŷ*^(var)^) (the baseline APARENT2 variant prediction)
- 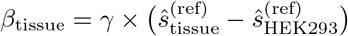 (the tissue-specific prediction)
- 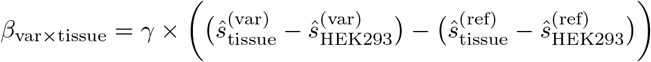 (the tissue-specific variant effect) and re-arrange the terms, we get:

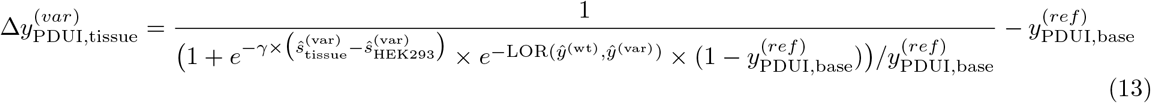

Compared to Equation 6, the difference is that we scale the variant odds ratio 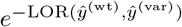predicted by APARENT2 with a tissue-specific odds ratio prediction 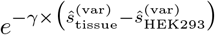. Note that there is a free hyper-parameter *γ ∈* ℝ that we need to tune on the GTEx aQTL data in order to properly scale the score residual 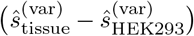 predicted by the tissue-model. We use the same value of *γ* for all tissue types (*γ* is chosen so as to maximize the median spearman r against measured 3’ aQTLs across all tissues).

### Mask-based Variant Interpretation

We adapted our recent work on mask-based interpretation [47] to find the contextual features within a sequence that explain the *relative* fold change between wildtype- and variant predictions. If the effect of all nucleotides were independent, the solution would simply be to return the mutated position itself (and nothing else). But, assuming mutations interfere with complex cis-regulatory code, the mask would have to retain a larger set of nucleotides (distant motifs, etc.) to reconstruct the variant effect. This is different from our earlier work, which focused on finding salient features that explain the *absolute* prediction of individual sequences. We found that per-example attribution worked stably for this task, so for simplicity we settled on optimizing individual masks rather than training a parametric ad-hoc interpreter and fine-tuning its scores. Let ***s*** *∈* (0, +*∞*]^*N*^ be the scores (the ‘mask’) that we will optimize specifically for the wildtype- and variant sequences ***x***^(wt)^ and ***x***^(var)^ of length *N*. We first set *s*_u_ = +*∞* and freeze this score, where *u* is the position of the mutation. Next, we channel-broadcast ***s*** into ***s***? *∈* (0, +*∞*]^*N×*4^ (same shape as ***x***^(wt)^):

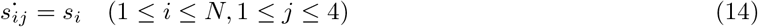

We then use 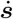 as interpolation coefficients between the original wildtype pattern ***x***^(wt)^ and a reference pattern 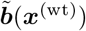 (taken here as a laplace-smoothed copy of the wildtype pattern 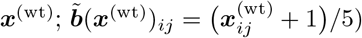:

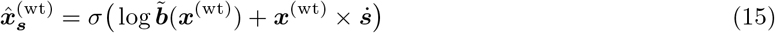

Here *σ* denotes position-wise softmax, i.e. 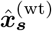is a softmax-relaxed position-specific scoring matrix (PSSM) whose entropy is controlled by ***s***. Next, we sample a discrete one-hot coded pattern 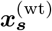from 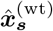using the Gumbel distribution [81]:

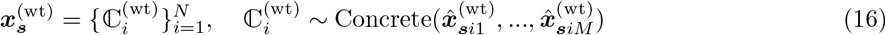

The next step is to construct a similar sample 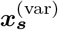of the variant pattern, whose information content has been masked and only the salient features marked by ***s*** are conserved. While we could theoretically re-apply Equation 15-16 to 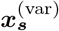 the same way we obtained 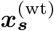, that approach does not work well in practice. The reason is that if the wildtype- and variant PSSMs 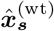 and 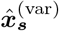have high entropy (which they are optimized for), then drawing independent samples from each PSSM will result in patterns with very different sequence content (except for the small set of features retained by ***s***). Consequently, the variance in the resulting predictions will be unnecessarily high. Instead, we directly construct the mutated sample 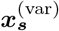 from the wildtype sample 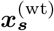 by ‘erasing’ the wildtype nucleotide and adding the mutation:

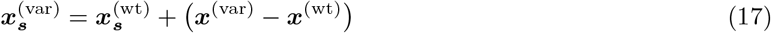

Both samples 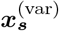 and 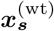 now have the same randomized (masked) background content and the same feature set retained by ***s***. Finally, we optimize the cost defined in Equation 18, which minimizes the mean squared error between the original and scrambled log odds ratio-predictions while maximizing entropy.

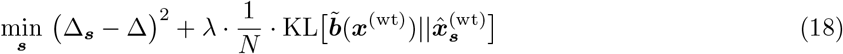

Here 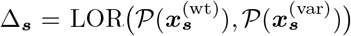 and Δ = LOR (*𝒫* (***x***^(wt)^), *𝒫* (***x***^(var)^)), where *𝒫* (***x***) is the proximal isoform proportion predicted by APARENT2 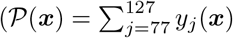 for predicted cleavage ***y***(***x***)). See Equation 5 for a definition of LOR. Note that we do not optimize ***s*** directly; instead we optimize parameters ***w*** *∈* ℝ^*N*^, which are instance-normalized and softplus-transformed into ***s*** (***s*** = Softplus(IN(***w***))). In our experiments, we optimize ***w*** for 300 iterations of gradient descent (Adam, learning rate = 0.01). We noticed more stable performance if we first optimize the mask ***s*** for a small target KL-divergence *t*_bits_ (Equation 19) for the first few gradient updates before maximizing the entropy unbounded.

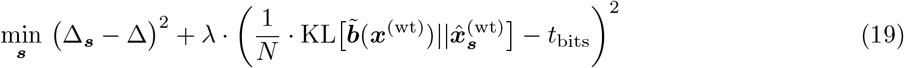

### ASD Cohort Data Filtering Procedure

The Autism Spectrum Disorder (ASD) WGS data from An et al. [49] was filtered by different criteria in some of the figures. In Supplementary Figure S5B (left), we remove variants (in cases and controls) that occur in a PAS which shares common variants in gnomAD (AF *>*0.01%) with strictly larger effect sizes or with *>*1.5-fold effect sizes. In Supplementary Figure S5B (right), we apply more stringent filtering (1.25-fold effect size cutoff) and we also remove variants that occur in PASs with a downstream neighboring PAS within 200nt in PolyA DB v3. In Main Figure 5G, we re-processed the gnomAD data by binning SNVs in the same PAS by their effect size (5 bins) and recomputed their (joint) allele frequency by aggregating allele counts in each bin. This allows for a group of rare variants (in the same PAS) to take the role of one common variant if their effect sizes are comparable. The same filtering procedure was used for the cohort data from Yuen et al. [73] (Supplementary Figure S5C), but the gnomAD AF cutoff was raised to 0.1% due to the smaller sample size. For both datasets, whenever a variant overlaps multiple PASs, we assign the mutation to the PAS with largest predicted effect size (we tested other assignment strategies in Supplementary Figure S5D).

## Availability of Data and Code

All code and data is available at http://www.github.com/johli/aparent-resnet. The variant prediction model is available online as an interactive web tool at https://apa.cs.washington.edu/. External software and data used in this study are listed in the Methods section.

## Acknowledgments

This work was supported by NIH award U01HG012069 to A.K. & NIH award 5R21HG010945 and NSF Award EF 2021552 to G.S.

## Author Contributions

J.L., A.K. and G.S. conceived and developed the project. J.L. performed the computational analyses. J.L., A.K. and G.S. wrote the paper.

## Supplemental Information

**Figure S1:**
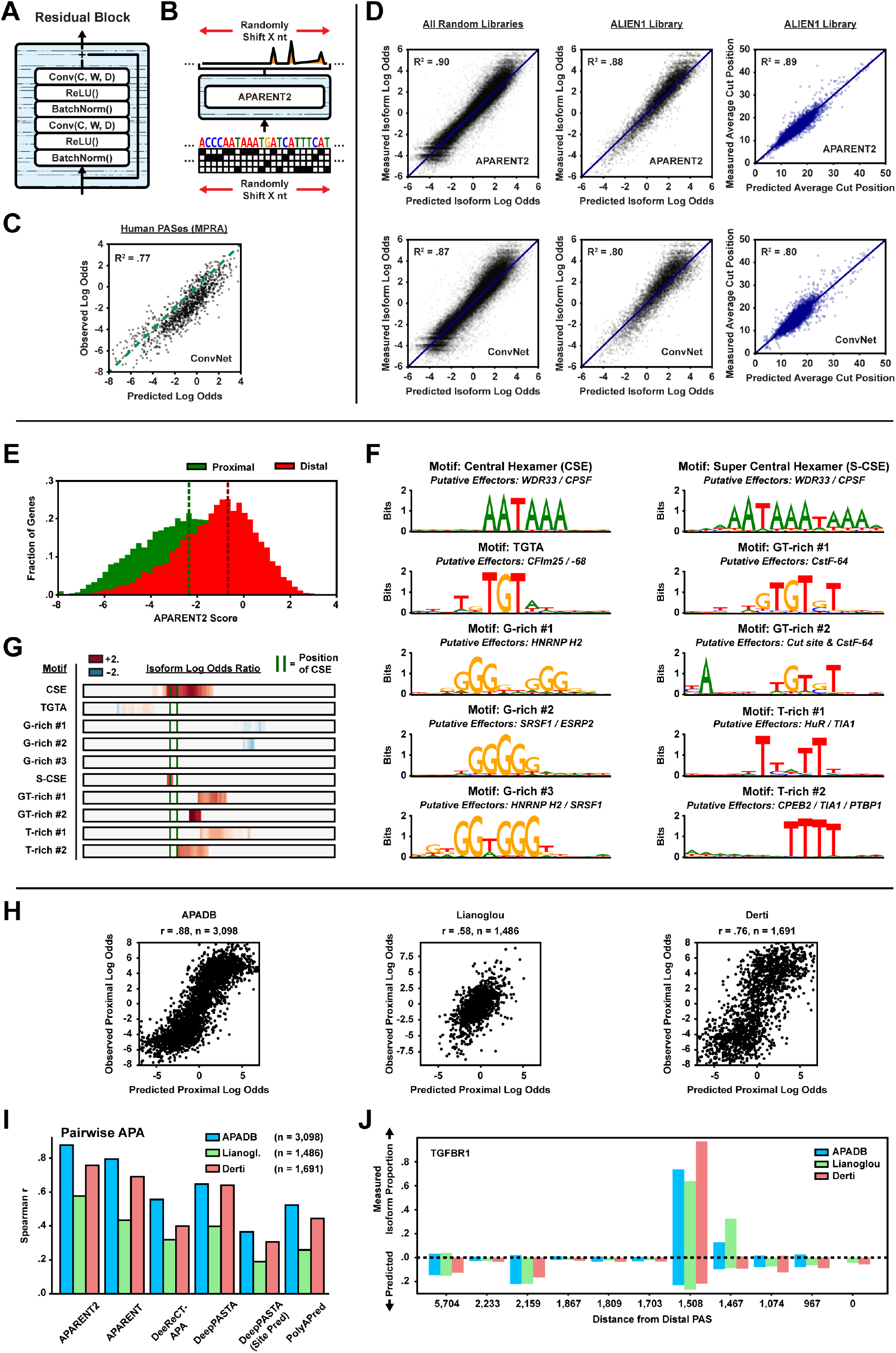
**A** The internal architecture of a residual block. C = # of channels, W = filter width, D = dilation rate. **B** During training, the input sequences and their target distributions are randomly shifted by a number of nucleotides. **C** Predicted vs measured isoform log odds of held-out human PASs measured in the MPRA of Bogard et al. [36] (*n* = 1, 085). **D** Predicted vs measured proximal isoform log odds of held-out test sequences from all MPRA libraries (left) (*n* = 60, 198) or the ALIEN1 library only (middle) (*n* = 7, 755), and predicted vs measured average cut position downstream of the CSE of ALIEN1 test sequences (right) (*n* = 6, 203). **E** Distribution of predicted isoform log odds using APARENT2 for the proximal-most and distal-most PASs in the 3’ UTR of *n* = 12, 503 genes. **F** Predicted vs observed isoform log odds between pairs of adjacent human 3’ UTR PASs, as measured in three separate native transcriptomic datasets [13, 41, 56]. Predictions are made by linear regression of the APARENT2 scores of the proximal and distal signals and their log-distance as features. Read counts were pooled across tissues. A minimum read count of 500 was used as cutoff for all three data sources. **G** Comparison of correlation between predicted and measured isoform log odds for pairs of adjacent human 3’ UTR PASs. Each model predicts logit scores which are used to fit a pairwise APA regressor (20-fold cross-validation). **H** Example multi-PAS isoform predictions and measurements for the gene TGFBR1. The predictions were made using the APARENT2 multi-PAS regressor (softmax regression with additive scores, no Saluki inputs). **I** Left: Selection of RNA binding protein (RBP) motifs generated by TF-MoDISco [54]. Each motif is represented as a position weight matrix (PWM). The motifs were generated from *n* = 20, 000 randomly sampled PAS sequences from PolyADB V3. Right: Estimated isoform log odds ratios in the presence of a given motif at a specific position in the sequence (i.e. the increase, *or* decrease, in predicted isoform log odds when a given motif is present).

**Figure S2:**
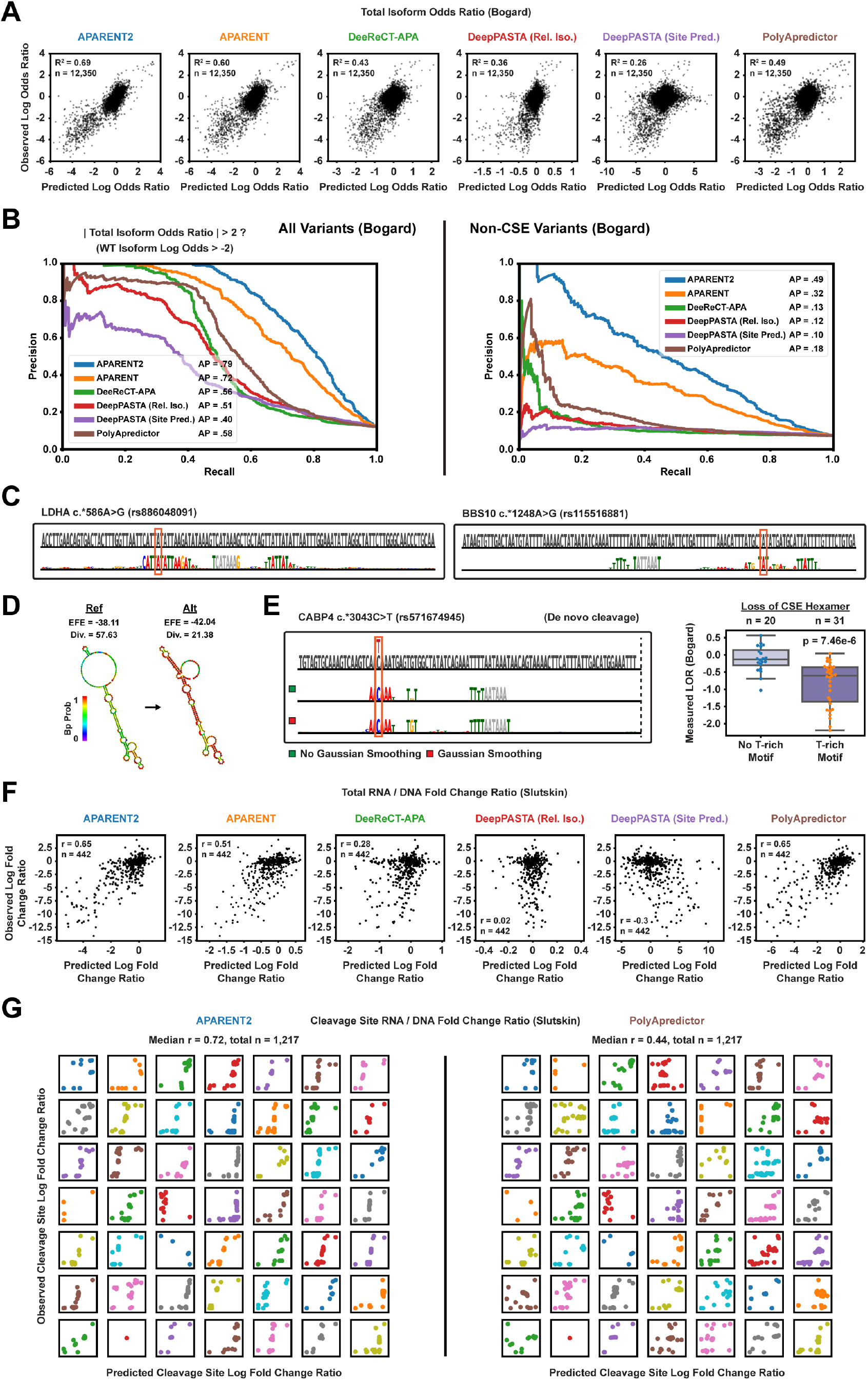
**A** Comparison of predicted vs measured variant isoform log odds ratios on the variant MPRA measured by Bogard et al. [36] (*n* = 12, 350). Individual scatter plots are shown for each model (APARENT2, APARENT, DeeReCT-APA, DeepPASTA Rel. Iso., DeepPASTA Site Pred. and PolyApredictor). **B** Comparison of precision-recall curves when tasking each of the models with classifying disruptive APA variants on the data from Bogard et al. [36]. Left: All variants. Right: Non-CSE variants only. The data only includes variants with a wildtype isoform log odds *> −*2. **C** Mask-based interpretation of example variants rs886048091 and rs115516881 with a fixed gaussian filter at the final scrambling layer (3 positions wide). **D** Secondary structure (thermodynamic free energy ensemble) of the wildtype and mutated PAS for variant rs886048091. Plots are produced with Vienna RNAFold [82]. EFE = Ensemble free energy, Div. = Ensemble diversity. **E** Interpretation of a ClinVar SNV that creates a de novo central hexamer element in the USE of an existing PAS (rs571674945). Two attributions are shown: An unregularized mask that is directly optimized to reconstruct the log odds ratio (LOR) (green) or a mask that has a gaussian filter hard-coded at the final layer (red). The boxplot shows measured log odds ratios from the MPRA of Bogard et al. [36]. The p-value is computed with a two-sided t-test. **F** Comparison of predicted vs. measured RNA/DNA log fold change ratios on the data from Slutskin et al. [35]. Individual scatter plots are shown for each model (APARENT2, APARENT, DeeReCT-APA, DeepPASTA Rel. Iso., DeepPASTA Site Pred. and PolyApredictor). The results are shown for variants of the PolyApredictor test set (*n* = 442). **G** Comparison of predicted vs. measured cleavage site RNA/DNA log fold change ratios on the data from Slutskin et al. [35]. Individual scatter plots are shown for each native wildtype UTR from the scanning mutagenesis experiment. Each data point corresponds to an individual wildtype cleavage position. Total *n* = 1, 217.

**Figure S3:**
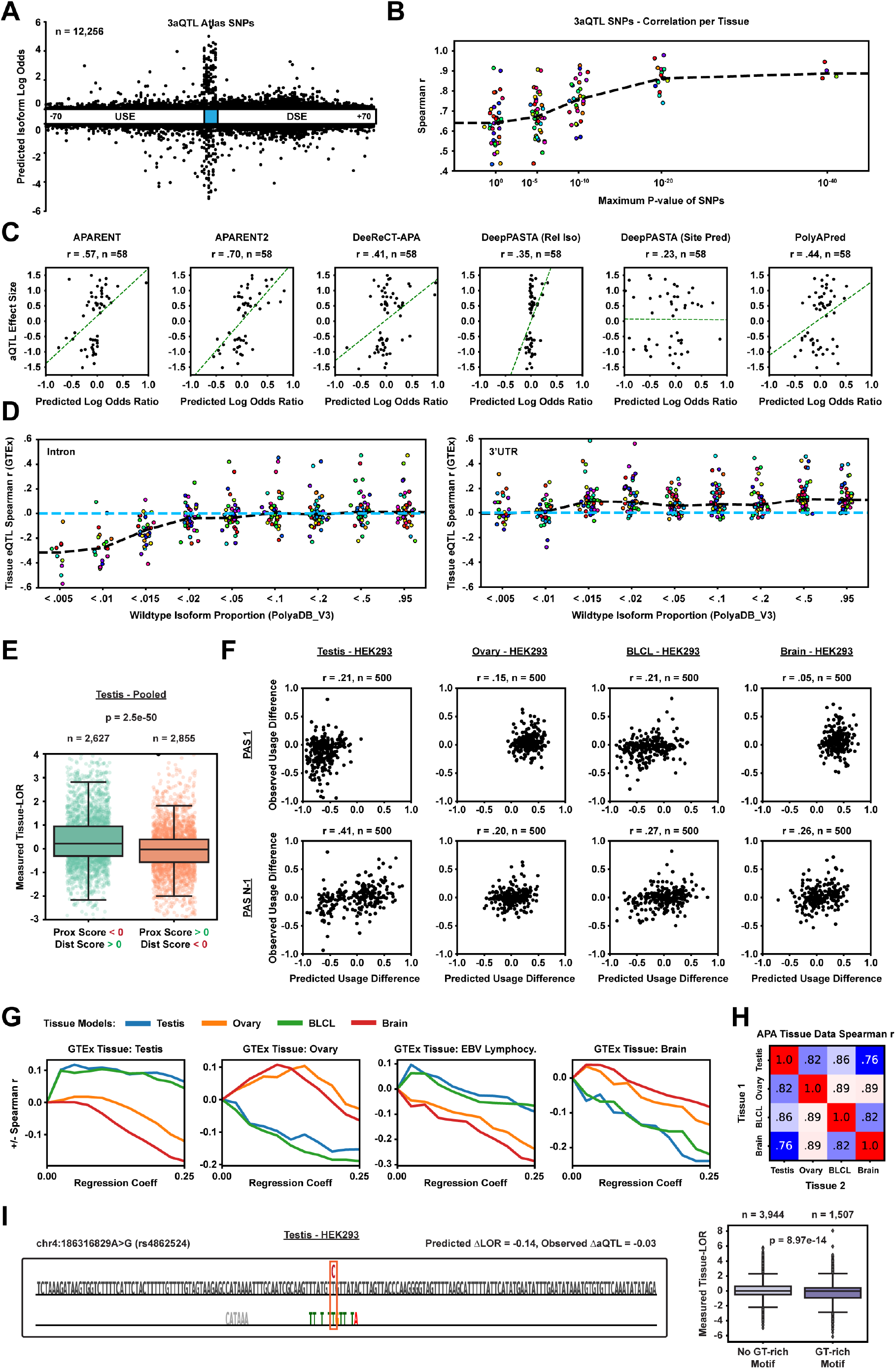
**A** Predicted isoform log odds ratio of SNPs from the GTEx v8 3’ aQTL atlas [78] overlapping annotated 3’UTR polyadenylation signals in PolyA DB v3 (*n* = 12, 256). The measured effect sizes of the GTEx v8 atlas are not publicly available, which is why the smaller v7 atlas is used in main Figure 3. **B** Correlation between predicted isoform log odds ratio (using APARENT2) and estimated 3’ aQTL effect sizes from the GTEx v7 atlas compiled by Li et al. [20]. Each dot corresponds to the spearman r correlation for a particular tissue, after having filtered the set of SNPs to those with a p-value below the cutoff specified by the x-axis. Number of unique lead cis-aQTLs across all tissues = 366. **C** Predicted isoform log odds ratio of each model vs estimated aQTL effect sizes of the Mittleman et al. [21] data (*n* = 58). **D** Correlation between APARENT2 isoform log odds ratios and estimated eQTL effect sizes for 1, 007 intronic GTEx eQTLs and 2, 225 3’ UTR eQTLs from Kerimov et al. [58], as a function of wildtype PAS usage as measured in tissue-pooled data from PolyA DB v3. **E** Measured difference in isoform log odds between testis and tissue-pooled data from Lianoglou et al. [41]. The left distribution is the subset of proximal PASs with APARENT2 scores *<*0 and distal APARENT2 scores (in the same gene) *>*0. Inversely, the right distribution is the subset of PASs with proximal APARENT2 scores *>*0 and distal scores *<*0. The p-value is computed with a two-sided t-test. **F** Predicted vs measured tissue-specific difference in isoform proportion between HEK293 and the target tissue, on a held-out test set of *n* = 500 genes from the Lianoglou et al. [41] data. Results are shown for either the most proximal, or next-to-last, PAS of each gene. **G** +*/−* Spearman r correlation w.r.t baseline APARENT2 predictions of GTEx aQTL effect sizes, separated by GTEx tissue type, as a function of regression coefficient *γ* (see Methods) for each tissue-specific model. **H** Correlation (sperman r) of PAS usage between tissues in data from Lianoglou et al. [41] (*n* = 6, 440 genes). **I** Reconstructive interpretation mask for a GTEx SNP (rs4862524) with a weak tissue-specific effect in Testis. Boxplot shows differential PAS usage (difference in isoform log odds) in data from Lianoglou et al. [41]. The p-value is computed with a two-sided t-test.

**Figure S4:**
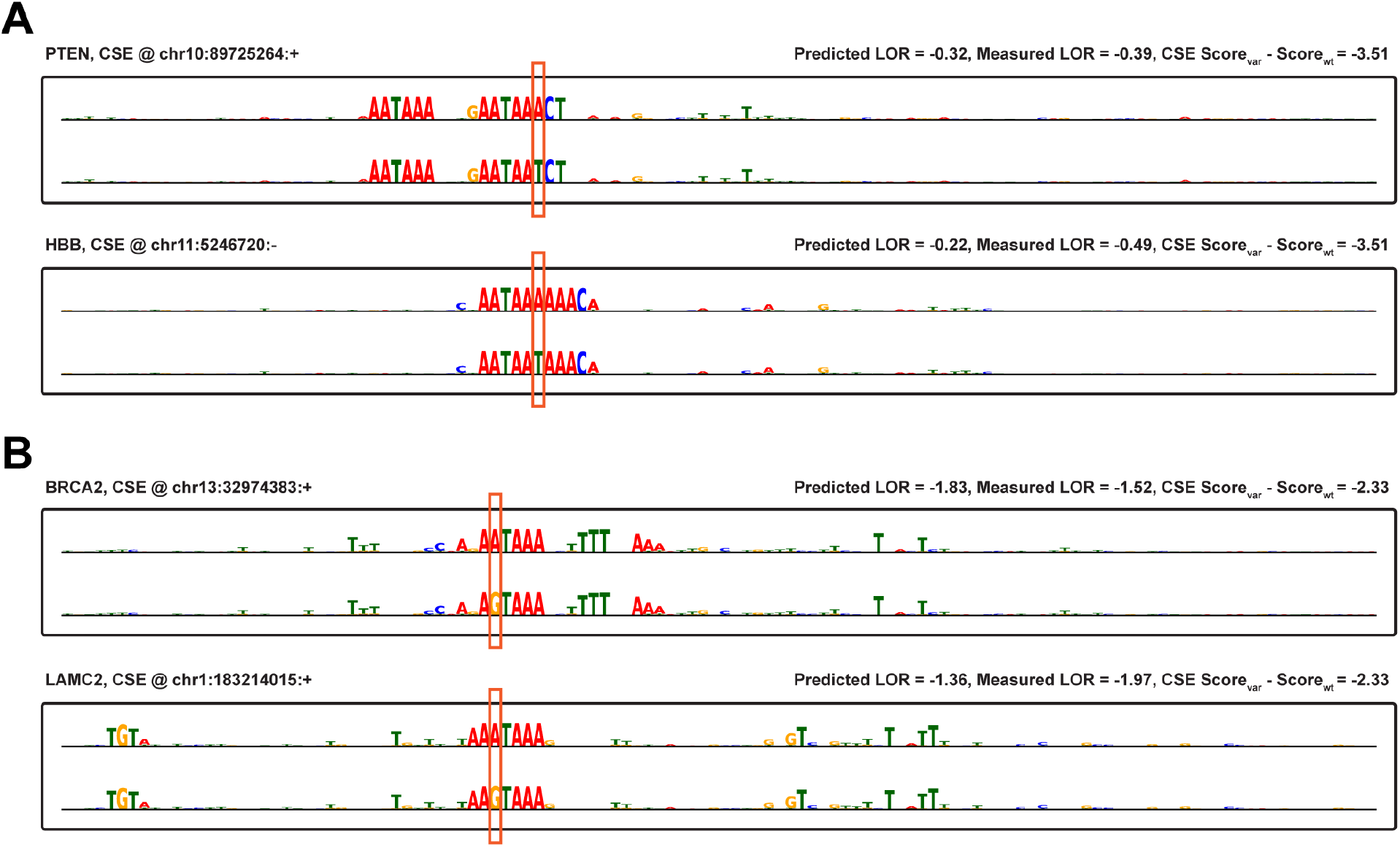
**A** Additional mask-based interpretations of functionally silent CSE mutations in the PTEN- and HBB genes. Annotated on the right are predicted and observed isoform log odds ratios (as measured in an MPRA), as well as CSE hexamer regression scores. **B** Additional interpretations of variants with dampened effect sizes (with respect to a linear hexamer regression model) in the BRCA2- and LAMC2 genes. Predicted and measured log odds ratios on the right.

**Figure S5:**
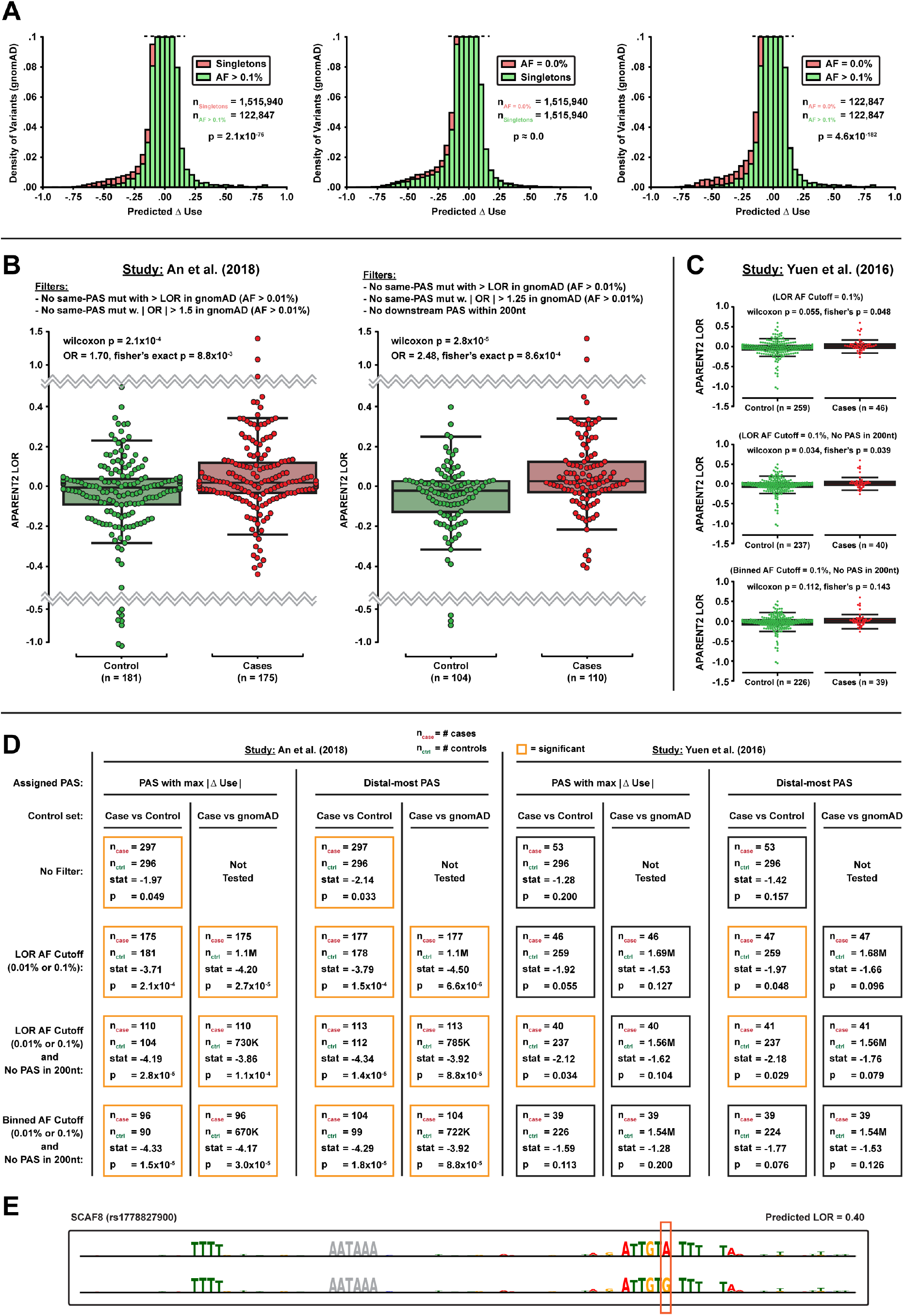
**A** Left: Predicted Δ isoform proportions for singletons (*n* = 1, 515, 940) and common variants (AF *>*0.1%; *n* = 122, 847) from gnomAD v3 [48] that overlap annotated PASs in PolyA DB V3. Middle: Comparison of predictions for a matched set of unobserved PAS variants (AF = 0.0%; *n* = 1, 515, 940) and singletons from gnomAD. Right: Comparison of predictions for a matched set of unobserved PAS variants (AF = 0.0%; *n* = 122, 847) and common variants (AF *>*0.1%) from gnomAD. **B** Distribution of predicted isoform log odds ratios among ASD cases and controls from the WGS study of An et al. [49]. Left: Case- and control variants are removed if they occur in PASs that have common variants in gnomAD (AF *>*0.01%) with larger effect size (log odds ratio) than the investigated variant or common variants that have an absolute odds ratio larger than 1.5 (*n*_control_ = 181, *n*_cases_ = 175). Right: Additional removal of variants that occur in PASs with a downstream PAS within 200nt in PolyA DB V3 (*n*_control_ = 104, *n*_cases_ = 110). A more stringent odds ratio threshold of 1.25 was used. This is the same filtering procedure as in Main Figure 5G, but here the allele count of variants within the same PAS with similar effect sizes have not been aggregated prior to filtering against gnomAD (see Methods for details on the variant filtering procedure). **C** Replicate analysis where the case variants are from the WGS study of 200 families from Yuen et al. [73] (the control variants come from An et al. [49]). The three filtering steps used in Supplementary Figure S5B and Main Figure 5G are also used here: (1) filtering variants with neighboring common mutations in gnomAD (AF *>*0.1%; *n*_control_ = 259, *n*_cases_ = 46), (2) additionally removing variants with downstream protective PASs within 200nt (*n*_control_ = 237, *n*_cases_ = 40), and (3) same filtering criteria as (2) but gnomAD AFs are re-calculated by aggregating allele counts of similar predicted effect size in the same PAS (*n*_control_ = 226, *n*_cases_ = 39). **D** Summary of statistical tests performed on the cohort data from An et al. [49] (left table) and Yuen et al. [73] (right table). P-values were calculated using Wilcoxon rank-sum tests. The rows denote different filtering criteria and the columns denote the control group used (controls either come from An et al. [49] or from an identically filtered view of gnomAD). Two different methods were tested for assigning variants to PASs (when variants overlap two nearby PASs): (1) the PAS with largest variant effect size or (2) the distal-most PAS. Significant tests are marked with an orange border. **E** Mask-based interpretation of an ASD-associated PAS mutation from Yuen et al. [73] (rs17778827900 in SCAF8).

**Figure S6:**
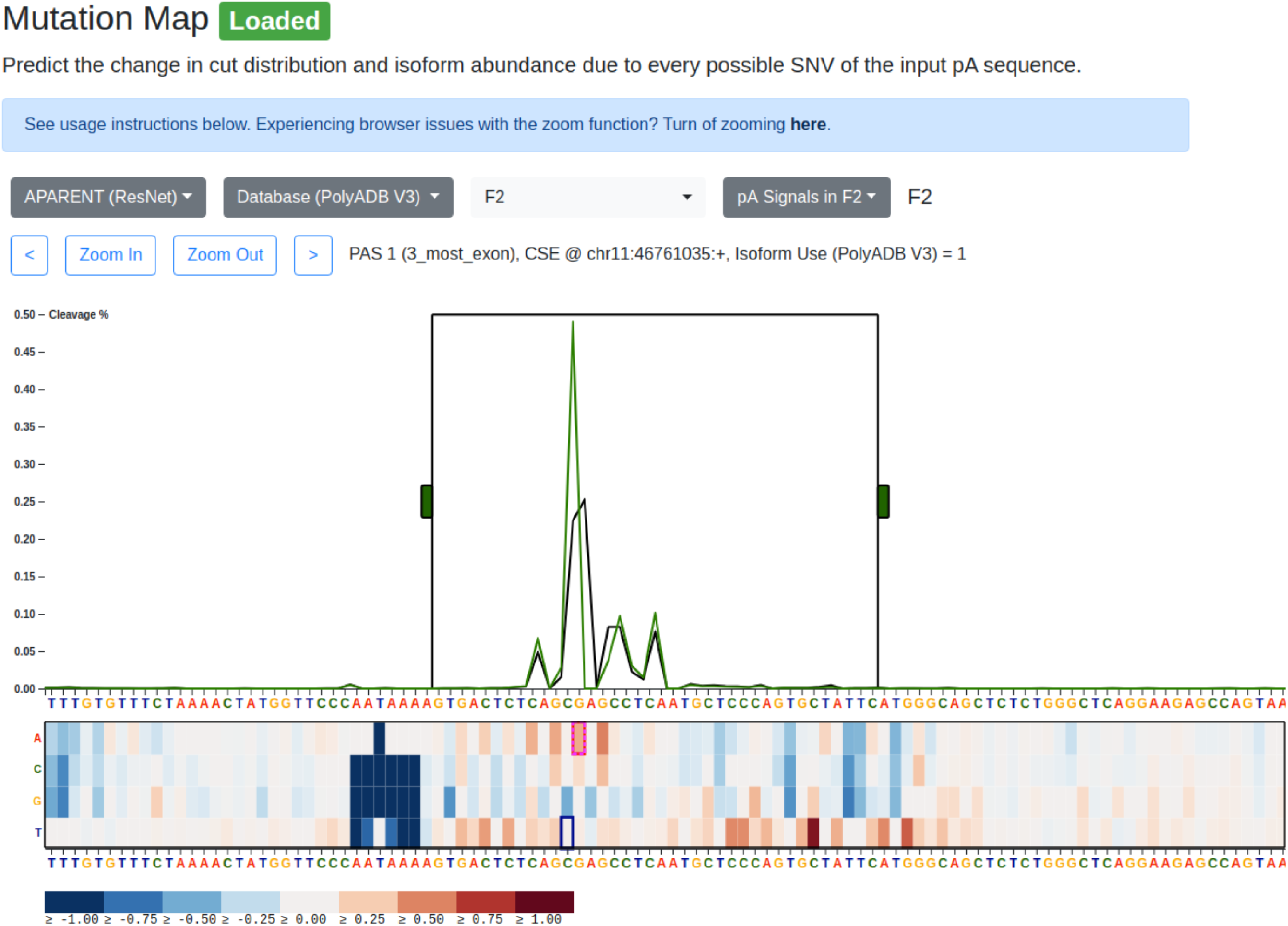
Screenshot of the polyadenylation variant prediction web tool. The tool allows users to perform in-silico saturation mutagenesis and interactively investigate the altered cleavage distribution due to every individual variant.

